# Clathrin differentially adapts its trimerisation domain during mammalian evolution to traffic the insulin-responsive GLUT4 glucose transporter

**DOI:** 10.64898/2026.08.20.745084

**Authors:** George T. Bates, Will P. Bultitude, Joshua Greig, Alyce McClellan, Nikos Pinotsis, Pallavi Ramsahye, Wingyan Skyla Siu, Kamila Kamuda, Riccardo Z. Chiozzi, Konstantinos Thalassinos, Snezana Djordjevic, Frances M. Brodsky

## Abstract

In humans, the CHC22 isoform of clathrin regulates glucose metabolism by trafficking the GLUT4 glucose transporter for intracellular storage in skeletal muscle and release following insulin signalling. Some vertebrate lineages have lost the gene encoding CHC22 but operate the same insulin-stimulated GLUT4 trafficking pathway. Here, we show that species lacking CHC22 exclusively produce an alternatively-spliced form of the universally expressed CHC17 clathrin isoform (CHC17-SAS) with a truncated C-terminus similar to CHC22, expressed predominantly in skeletal muscle. Through its trimerisation domain, CHC17-SAS binds the CHC22-specific adaptor SNX5 that enables CHC22’s distinct intracellular function. The 2.3 Å crystal structure of the CHC22 trimerisation domain demonstrates conservation of the core trimeric fold from CHC17 but differences in electrostatic surface charge that may account for their differential properties. Using GLUT4 translocation assays in HeLa cell models, we show that CHC17-SAS is a functional surrogate for CHC22. Identification of CHC17-SAS resolves the evolutionary conundrum posed by CHC22 absence in some vertebrate lineages, and reveals a common mechanism for mammalian GLUT4 trafficking.

## Introduction

Glucose metabolism is tightly regulated to ensure energy homeostasis and its dysregulation is a hallmark of metabolic disease. Skeletal muscle is responsible for 70% of insulin-stimulated glucose uptake^1,2^, which at the cellular level, is largely regulated by membrane traffic of the glucose transporter GLUT4^3^. In basal conditions, where no significant glucose uptake is required, GLUT4 is transported to an intracellular storage compartment, known as the GLUT4 storage compartment (GSC). This sequesters GLUT4 away from the plasma membrane, preventing it from importing glucose, while acting as a primed pool that can be rapidly mobilised to the cell surface upon insulin stimulation. In humans, this GLUT4 trafficking pathway is mediated through the action of two clathrin heavy chain (CHC) isoforms – CHC17 and CHC22. CHC22 captures GLUT4 from the ER-Golgi intermediate compartment (ERGIC) following its synthesis, bypassing the Golgi and transporting it to the GSC^4–6^. Once GLUT4 is released to the cell surface by insulin signalling, it is recaptured by CHC17-mediated endocytosis and sorted via endosomes and the trans-Golgi network (TGN) back to the GSC in a CHC22-dependent step^7^. As a result of its essential role in human GLUT4 traffic, siRNA-mediated knockdown of CHC22 is sufficient to cause the dissolution of the GSC in a HeLa cell line model^4^ and in human myotubes^5^. The clathrin heavy chain isoforms, CHC17 and CHC22, share 85% sequence identity and operate using the same general membrane transport mechanism, involving membrane recruitment, membrane deformation via self-assembly, and scission to form a clathrin-coated vesicle (CCV) to capture cargo. However, as exemplified by their roles in GLUT4 trafficking, CHC17 and CHC22 have distinct functions^8^. CHC17 carries out a range of housekeeping roles including clathrin-mediated endocytosis, protein sorting at endosomes and the TGN, while the only identified function for CHC22 is to traffic GLUT4 to the GSC. The tissue expression of CHC22 mirrors that of GLUT4 in muscle, while CHC17 is ubiquitously expressed. Additionally, CHC17 is expressed in all eukaryotes, while the *CLTCL1* gene encoding CHC22 is missing (or a pseudogene) in several vertebrate lineages including mice and rats, and ruminants, such as cows and sheep^9,10^. The convergent evolutionary losses of CHC22 are notable in the context of glucose metabolism, as many species lacking CHC22 (e.g. mice) traffic GLUT4 into an insulin-sensitive GSC similarly to humans, despite the absence of the protein that is required for the pathway in humans. The study reported here addresses this evolutionary conundrum through further evolutionary and structural characterisation of clathrin heavy chains.

The adaptors that recruit CHC17 to intracellular membranes are well characterised and recent studies have elucidated the molecular mechanism distinguishing CHC22 localisation from CHC17, enabling its specialised function^4,6,11^. Specific to CHC22 and not CHC17 is a bipartite interaction with the ERGIC tether p115, directly via the N-terminal domain of CHC22, and through the sorting nexins SNX5 or SNX6, which bind to CHC22 at its C-terminus and additionally link CHC22 to p115^4,6^. The tether p115 complexes with IRAP, which in turn links the trafficking machinery to the GLUT4 cargo^4,12^. Following the insulin-stimulated release of GLUT4 to the plasma membrane and CHC17-dependent re-uptake, GLUT4 is recycled back to the GSC from endosomes via the TGN^7^ by CHC22 interacting with the AP1 adaptor (also a partner of CHC17) plus sortilin in a complex distinct from CHC22’s secretory pathway interactions. It has been proposed that with the lack of CHC22 to traffic GLUT4 from the ERGIC to the GSC, rodents primarily rely on the endocytic-retrograde pathway to package GLUT4. However, rodent studies have shown that GLUT4 transits to the GSC before sampling the cell surface^13,14^ and formation of the GSC is dependent on ERGIC tether p115^12^, suggesting a potential CHC22-independent biosynthetic route of GLUT4 trafficking into the GSC. However, the molecular mechanism by which rodents compensate for the absence of CHC22 from their GLUT4 trafficking pathway remains unresolved.

Alternative splicing is a prevalent mechanism to modulate cellular behaviour with genetic diversity expanded through the regulated inclusion or exclusion of exons. In membrane trafficking, alternative splicing is often used to generate tissue-specific variants that fine-tune pathways for tissue-specific requirements^15^. For example, neuron-specific splicing of the regulatory clathrin light chain (CLC) subunit allows CHC17 to more efficiently carry out synaptic vesicle recycling in neurons^16^. We therefore addressed whether, instead of maintaining duplicated genes encoding two distinct protein isoforms (CHC17/CHC22) for GLUT4 trafficking, as is the case for humans, rodent and ruminant species could achieve a similar outcome by alternative splice variants of CHC17. Recent studies have shown that the regulated inclusion of exon 31 (residues VDAIKEK) into CHC17 modulates force production in murine muscle by altering its self-assembly properties^17^. Exon 31 inclusion alters the CHC17 sequence within the domain that mediates clathrin trimerisation, which is essential for its self-assembly properties and trimer stability and is known to be fine-tuned by CLC binding^18^. Furthermore, the CHC17 trimerisation domain (TxD) facilitates clathrin uncoating from clathrin-coated vesicles via a “QLMLT” motif that interacts with uncoating components Hsc70 and auxilin^19–21^. As the C-terminal sequence of the CHC22 TxD is shorter by 35 residues compared to CHC17, lacks this uncoating motif, and is the domain that binds SNX5^6^, we also investigated the relative properties of CHC TxDs as potential mediators of functional specialisation.

Here, we demonstrate that species lacking a gene encoding CHC22 express an alternatively spliced variant of CHC17 (CHC17-SAS) that functionally mimics CHC22 by truncation of its C-terminal TxD. The presence of CHC17-SAS or CHC22 in the 18 mammalian species analyzed is mutually exclusive and each are primarily expressed in muscle tissue with similar expression levels relative to CHC17. Biochemical characterisation of full length CHC17 (CHC17-FL), CHC17-SAS and the previously-identified CHC17-VDAIKEK splice variant, shows that alternative splicing modulates clathrin trimer stability as well as protein-protein interactions including SNX5 binding. Through structural analysis of the crystallised CHC22 TxD at 2.3 Å resolution, we present the first experimentally determined structure of CHC22, providing further insight into clathrin heavy chain architecture and protein interactions. Finally, we demonstrate the functional equivalence of CHC22 and CHC17-SAS in insulin-stimulated GLUT4 membrane traffic using an established HeLa cell model^4,6,22^.

## Results

### Loss of *CLTCL1* is correlated with the expression of an alternatively spliced variant of *CLTC*

An expanded analysis of the presence of gene *CLTCL1* (encoding CHC22) was performed using a set of 121 mammalian genomes to investigate its evolutionary conservation. These genomes were stringently filtered to minimize the risk of mischaracterizing gene absence from incomplete assemblies. As expected, regions homologous to *CLTC* (encoding CHC17) were identified in all species. Upon searching for *CLTCL1*, homologous regions were only identified in 67 species, implying that *CLTCL1* has been lost in 54 of the 121 assessed mammalian species (Supplementary Table 1).

A phylogenetic tree was constructed to assess the clade-specific losses of *CLTCL1* (Fig. 1) and indicated that *CLTCL1* has been lost in more mammalian lineages than had been previously identified^10^. *CLTCL1* was absent in all 32 analyzed members of the Cetartiodactyla order, including ruminants (all available), pigs (*S. scrofa*) and whales (*G. melas*), and in the single available Monotremata species. *CLTCL1* displays a mixed presence in the Rodentia order, absent in 17 of the 20 species, including mouse (*M. musculus*) and rat (*R. norvegicus*), but present in the Sciuridae sub-family (e.g. grey squirrel; *S. carolinensis*). Similarly, the presence of *CLTCL1* in the Chiroptera order of bats appears to be split between the old-world Yinpterochiroptera (e.g. *P. giganteus*) which retained *CLTCL1*, and the new-world Yangochiroptera (e.g. *P. hastatus*) which have lost it. While the losses of *CLTCL1* in several model species, e.g. mouse and cow, have been reported previously^9,10^, this analysis with a greatly expanded dataset further validates these conclusions, demonstrating that these species are representative within their evolutionary cohort.

**Figure 1.**
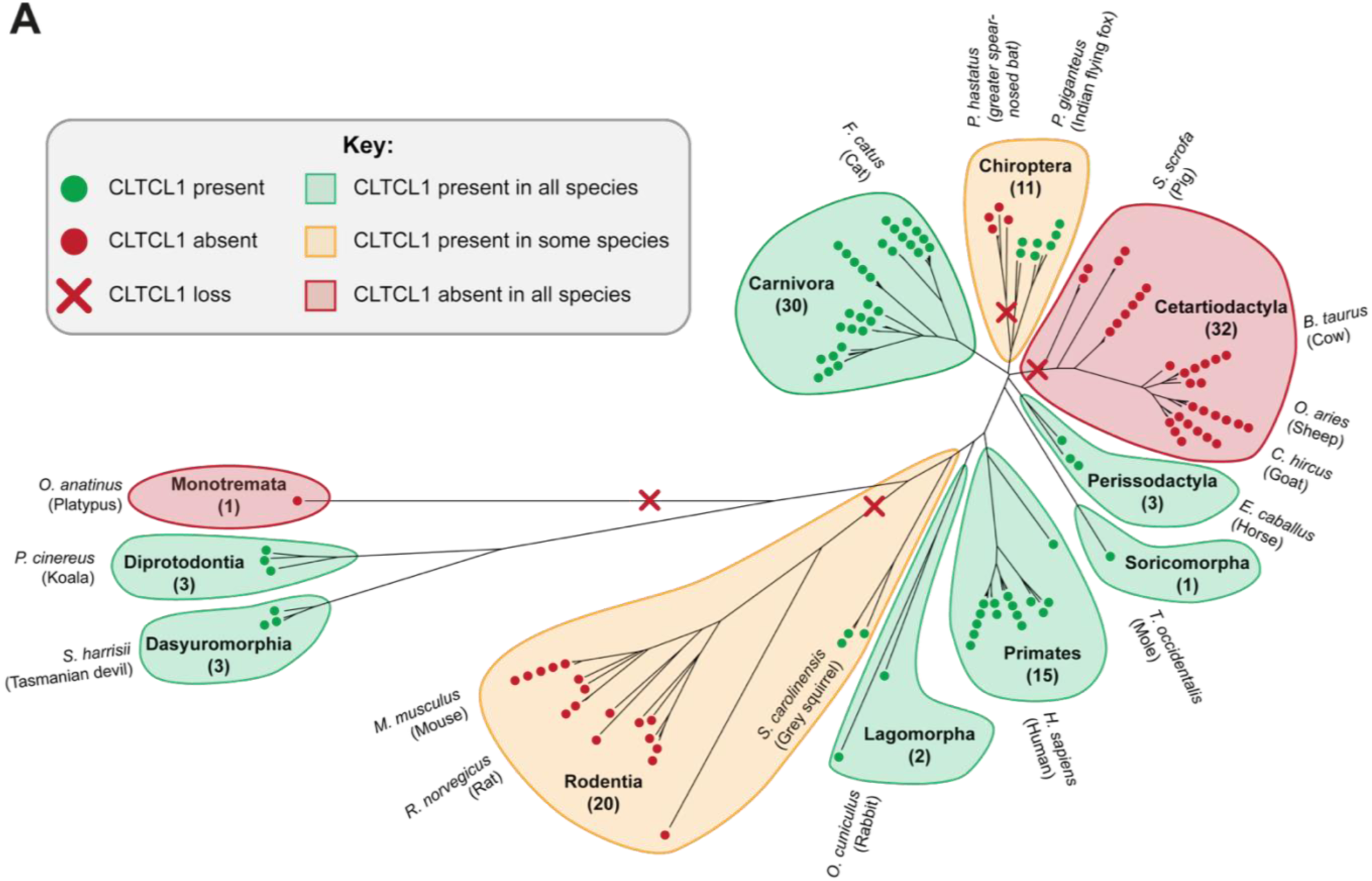
Expanded mammalian phylogenetic analysis reveals the widespread losses of *CLTCL1* (CHC22) within the Rodentia and Cetartiodactyla orders. **(A)** A phylogenetic tree, based on *CLTC* (encoding CHC17) sequences, from a filtered set of high-quality 121 mammalian genomes annotated to depict the presence or absence of *CLTCL1* (encoding CHC22). The number of genomes within each of the 11 represented mammalian orders are denoted in brackets and colored circles with red or green indicating absence or presence of CLTCL1. The orders are colored green, orange or red depending on the representation of species encoding *CLTCL1*, with one or more representative species labelled for clarity.

To investigate whether an alternatively spliced version of *CLTC* may compensate for the loss of *CLTCL1*, the skeletal muscle transcriptome was examined within *CLTCL1*-expressing and non-expressing mammalian species mining publicly available databases (Supplementary Table 1). Where existing transcriptomic data permitted analysis, species lacking *CLTCL1* were found to express a shortened transcript of *CLTC* that utilizes an alternative final exon (Fig. 2A), hereby termed short alternatively spliced (SAS). This produces a truncated protein, where the final 41 residues of CHC17 are replaced with 10 or 11 residues from the alternate final exon in Rodentia and Cetartiodactyla, respectively (Fig. 2B). While the genomic location of the alternate final exons and their encoded residues differ between the Rodentia and Cetartiodactyla orders, this alteration mirrors the sequence of CHC22, with a shorter C-terminus relative to CHC17 and 6 residues of unshared sequence appended.

**Figure 2.**
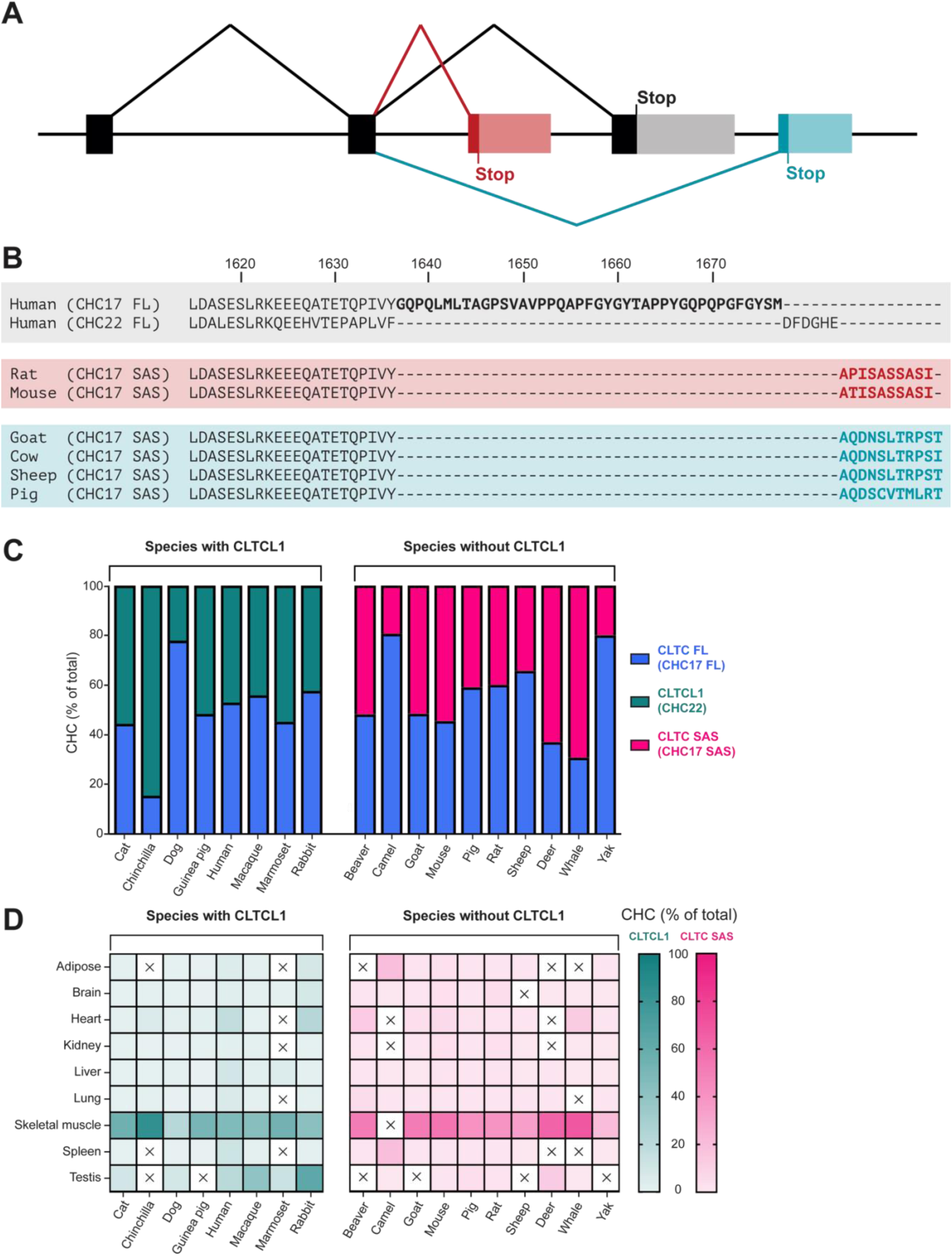
Species that lack *CLTCL1* encode an alternatively spliced *CLTC* transcript that resembles *CLTCL1* in skeletal muscle. **(A)** A schematic depicting the splicing of the final exons of *CLTC*. Black exons represent those found in conventional *CLTC*, red and blue exons represent the alternatively spliced final exons from Rodentia and Cetartiodactyla, respectively. Lighter shades represent the 3’ UTR. Splicing events denoted above and below the exons by solid lines. **(B)** Sequence alignment of the C-terminus of different CHC protein variants (1613-end) identified from transcriptome analysis. Sequences are shaded grey, red and blue for human, rodent and Cetartiodactyla proteins, respectively. FL refers to full-length, conventional CHC, SAS refers to the short alternatively spliced CHC. **(C)** The transcript abundance of *CLTCL1*, *CLTC-FL* and *CLTC-SAS*, expressed as a % of the total of these three transcripts, in CLTCL1-expressing and non-expressing mammalian species. Transcript abundance calculated using exon junction counts. **(D)** The transcript abundance of *CLTCL1, CLTC-FL* and *CLTC-SAS*, expressed as a % of the total of these three transcripts, from different tissues. X indicates data are unavailable. Transcript abundance was calculated using exon junction counts.

### *CLTCL1* and *CLTC-SAS* are expressed in a mutually exclusive manner in skeletal muscle

Notably, the transcriptomic analysis here suggested that the *CLTC-SAS* transcripts were only detected in species that lacked *CLTCL1*, implying that the expression of CHC22 and CHC17 SAS is mutually exclusive. To validate this observation and to provide a semi-quantitative analysis of *CLTC-SAS* expression, the relative expression levels of each clathrin transcript subtype were examined from skeletal muscle using exon junction counts (Fig. 2C). Species that express *CLTCL1* were found to contain no *CLTC-SAS* transcripts, and species that have lost *CLTCL1* expressed *CLTC-SAS* but no *CLTCL1* (converted to a pseudogene in some cases^9^). Moreover, across the analyzed species, the mean transcript abundances of *CLTCL1* (50.3%) and *CLTC-SAS* (44.5%) were similar relative to full length *CLTC* (*CLTC-FL)*. Broadening the scope of the analysis to include a range of additional tissues showed that *CLTC-SAS,* just like *CLTCL1*, is expressed most highly in skeletal muscle (Fig. 2D), with limited expression elsewhere.

To verify that the *CLTC-SAS* transcript is translated and that the CHC17-SAS protein is expressed, protein lysates were examined from mouse (*Mus musculus*), as a representative model of a *CLTC-SAS*-expressing species. CHC17 was immunoprecipitated from skeletal muscle and kidney, tissues that express the transcript in high and low abundances, respectively. Analyzing the immunoprecipitates by Coomassie-stained SDS-PAGE gel showed a doublet visible from skeletal muscle tissue, while only a single band was present from kidney tissue (Fig. 3A). This is consistent with skeletal muscle expressing two distinct CHC17 variants: full length (CHC17-FL) and CHC17-SAS. In agreement with the exon junction counts (Fig. 2C), quantifying the band intensities showed that in skeletal muscle, the two CHC17 isoforms appear to be present at roughly equal levels (Fig. 3B). Mass spectrometry sequencing of the immunoprecipitated bands further confirmed that the doublet is composed of CHC17-SAS and CHC17-FL in skeletal muscle, while the equivalent region in kidney only contains CHC17-FL (Fig. 3C, D).

**Figure 3.**
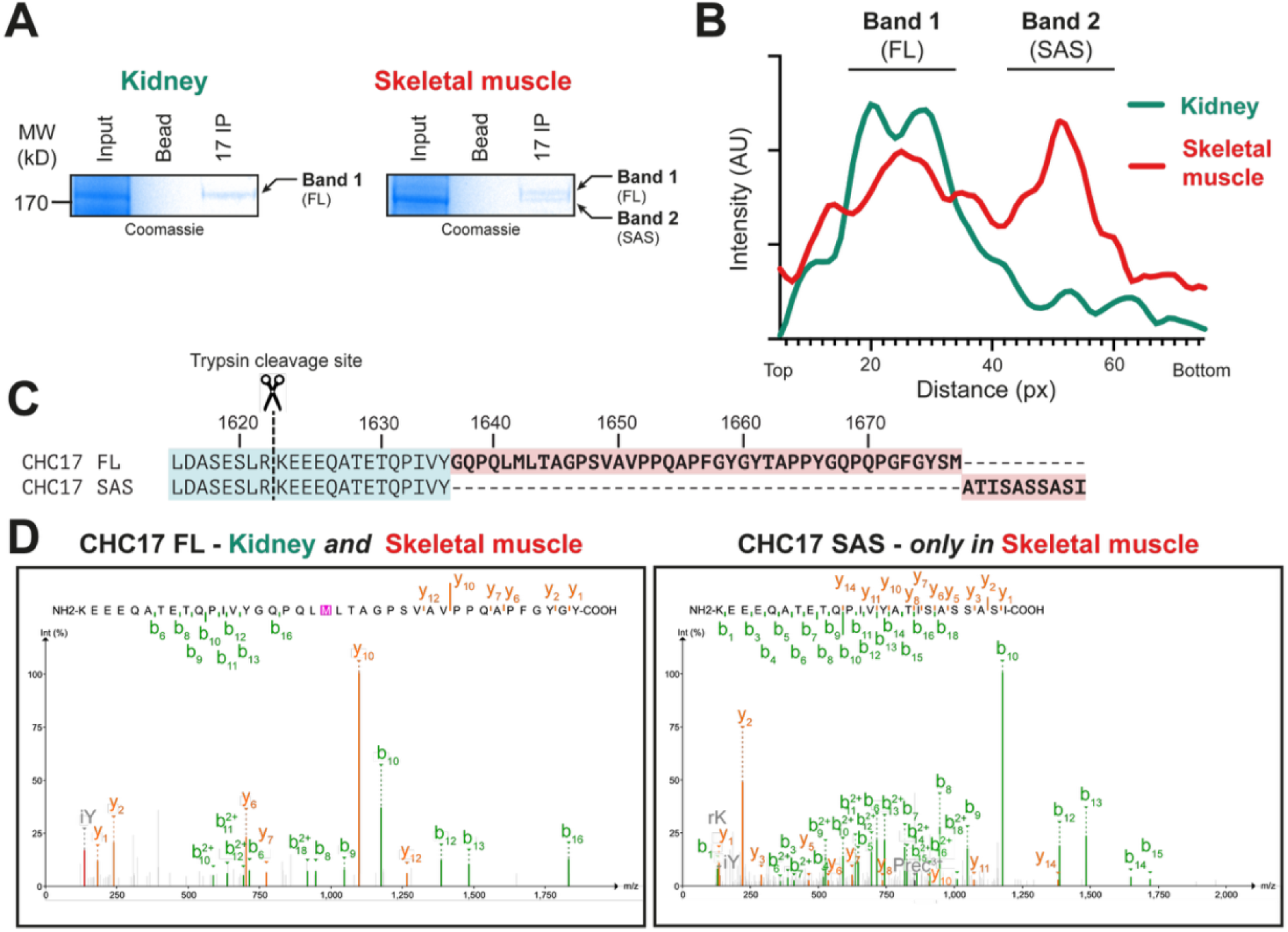
CHC17-SAS is present in mouse muscle tissue. **(A)** Immunoprecipitates of CHC17 from mouse kidney or skeletal muscle tissue visualised on a Coomassie-stained SDS-PAGE gel. Lanes show Input (5%), Bead (no antibody control) and 17 IP (CHC17 immunoprecipitate). Position of molecular weight markers is indicated in kD. **(B)** Quantification of the CHC17 immunoprecipitates in **(A)**. An intensity profile was plotted across the CHC17 containing sections of the 17IP lanes from kidney (green) and skeletal muscle (orange) **(C)** Sequence alignment of CHC17-FL and CHC17-SAS, showing the predicted peptide generated from trypsin cleavage and the residues that are shared (blue) or divergent (red). **(D)** Tandem mass spectrometry spectra of peptides detected in bands excised from the immunoprecipitates shown in **(A)**. The b-ions (green) and y-ions (orange) refer to fragments that include the N- or C-terminus of the peptide, respectively. The numbers indicate the number of residues included in each fragment.

Together, this data demonstrates that the species expression of *CLTCL1* and *CLTC-SAS* transcripts is mutually exclusive, but where they are expressed, they are present at similar levels to each other and share similar tissue-specific expression patterns. Moreover, the *CLTC-SAS* transcript is translated into protein and can be detected in mouse skeletal muscle, where it is present in approximately equal amounts to *CLTC-FL*.

### Short alternative splicing maintains trimer stability despite structural differences

Having validated the existence of CHC17-SAS, the impact of splicing on the biophysical properties of CHC17 was then characterized. The stability of the trimer formed by CHC17-FL was compared to the stability of the CHC17-SAS trimer, derived from the murine sequence, and to the other previously characterised alternatively-spliced human variant CHC17-VDAIKEK, which also alters the TxD^17^ (Fig. 4A).

**Figure 4.**
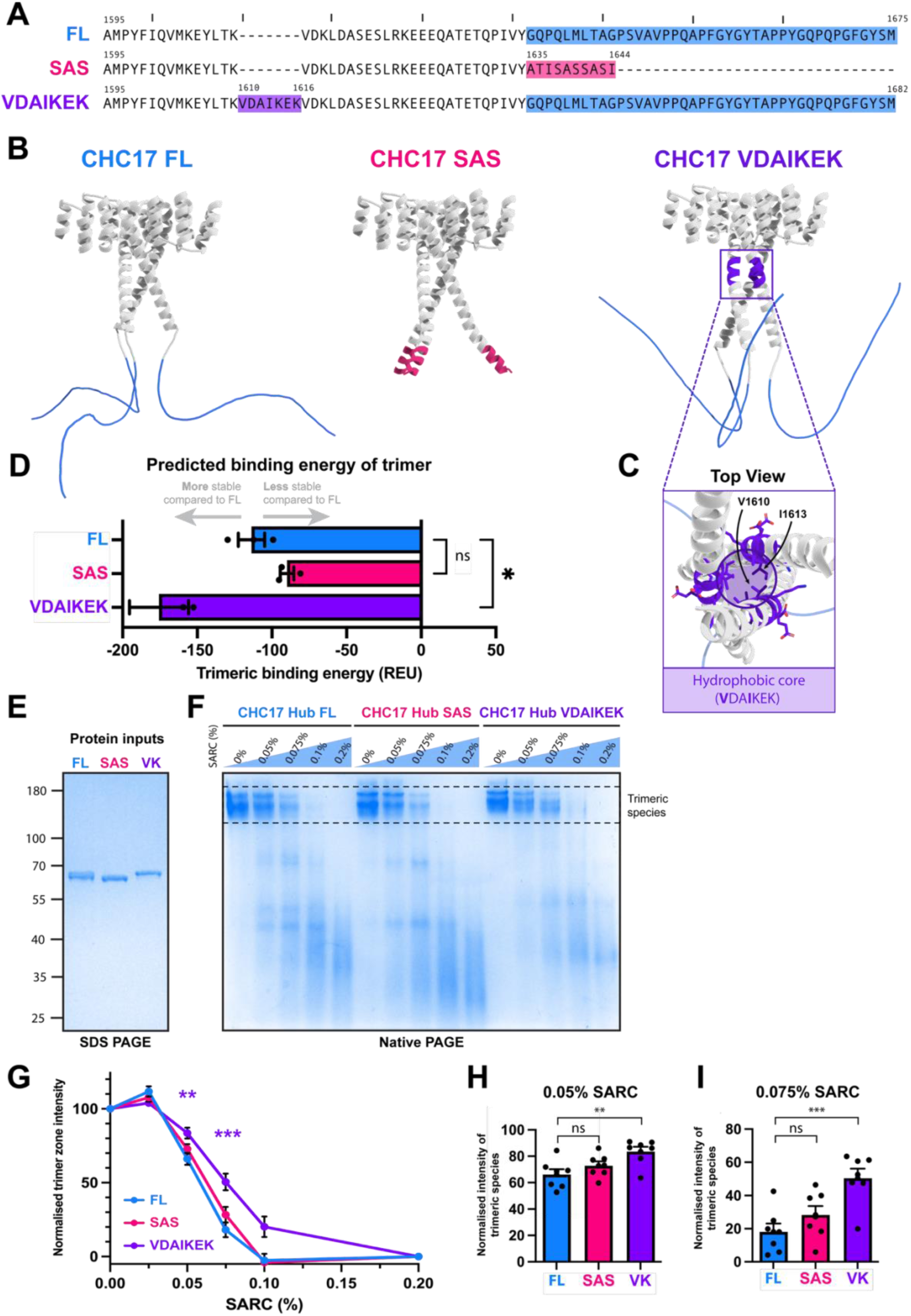
Structural and biochemical analysis reveals differential effects of alternate splicing on trimeric stability. **(A)** Sequence alignment of the CHC17-FL, CHC17-SAS and CHC17-VDAIKEK C-termini (residues 1595-end). Residues encoded by the conventional final exon are shaded in blue, the alternative final exon in pink and Exon 31, encoding VDAIKEK, in purple. **(B)** AlphaFold-Multimer models of CHC17-FL, CHC17-SAS and CHC17-VDAIKEK TxDs (residues 1522-end). Residues colored as described in **(A)**. **(C)** Inset from the boxed area of **(B)** showing a top-down view of the trimerisation interface of CHC17-VDAIKEK. **(D)** Predicted binding energy of the trimeric interface from the CHC17-FL, CHC17-SAS and CHC17-VDAIKEK AlphaFold models, calculated using Rosetta InterfaceAnalyzer and expressed in Rosetta Energy Units (REU). Error bars display standard error from three independent relaxation trajectories per construct. **(E)** SDS-PAGE gel displaying the purified Hub regions of CHC17-FL, CHC17-SAS and CHC17-VDAIKEK **(F)** Native PAGE gel of the detergent challenge assay. CHC17-FL, CHC17-SAS or CHC17-VDAIKEK hubs were incubated with 0%, 0.05%, 0.075%, 0.1% or 0.2% SARC detergent. Dashed box indicates the region of the trimeric species used for quantifications in **(G)** – **(I)**. **(G)** Quantification of detergent challenge assay in **(F)**, measuring the band intensity within the “trimer zone” as a % of the intensity of 0% SARC. Statistical analysis was performed using a one-way ANOVA and represents significance between FL and VDAIKEK conditions, as is also annotated in (H)/(I). ns = not significant, *P < 0.05; **P < 0.01; ***P < 0.001. **(H) /(I)** Subset of data shown in **(G)** for 0.05% and 0.075% SARC (n=7). Error bars represent standard error of mean. ns = not significant, *P < 0.05; **P < 0.01; ***P < 0.001

The TxDs of CHC17-FL, murine CHC17-SAS and human CHC17-VDAIKEK were modelled as trimers using AlphaFold-Multimer^23,24^ to predict the structural impact of the alternative splicing (Fig. 4B). As expected, the CHC17-FL predicted model aligns closely with previously solved experimental structures^25–27^, yielding an RMSD of 1.18 Å over 312 residues (Supplementary Fig. 1; PDB: 6SCT) and trimerization interactions attributed to the tripod helix. The CHC17 residues encoded by the conventional final exon of *CLTC* are typically absent from structures due to their flexibility^26,27^. Consistent with these observations, the models predict these residues to be disordered and lack secondary structure (Fig. 4B). Modelling CHC17-SAS shows that the splicing removes these disordered residues, with the replacement residues extending the previous α-helix outwards, away from the trimerisation interface. In contrast, in CHC17-VDAIKEK, the additional residues from the alternative splicing are inserted within the trimeric interface, with residues V1610 and I1613 projecting inwards to form a stabilising hydrophobic core (Fig. 4C). Using these three AlphaFold models, the interfacial binding energy was then calculated to predict clathrin trimer stability (Fig. 4D). This modelling suggested that the CHC17-VDAIKEK trimer is more stable than CHC17-FL due to the increased interfacial surface area, while the CHC17-FL and CHC17-SAS TxDs are predicted to be similar in stability due to their differences being observed in the disordered segment beyond the trimerisation interface.

To experimentally validate the above *in silico* analysis, clathrin trimer stability was assessed biochemically using a detergent challenge assay^18^. The Hub regions (residues 1074 to the C-terminus, comprising the TxD plus the proximal leg) of CHC17-FL, murine CHC17-SAS and human CHC17-VDAIKEK were recombinantly expressed and purified (Fig. 4E), followed by incubation with increasing concentrations of the detergent SARC to dissociate trimers into monomers (Fig. 4F). Quantifying the trimerized Hub levels using Native-PAGE showed that while all three Hub variants are fully trimeric in the absence of SARC, CHC17-VDAIKEK dissociated into monomers to a lesser extent than CHC17-FL, with the biggest differences seen at 0.05% and 0.075% SARC (Fig. 4G-I). As predicted *in silico*, CHC17-SAS Hub showed no significant difference in trimer stability relative to CHC17-FL Hub. This analysis demonstrates that alternative splicing can affect the CHC17 TxD, resulting in structural rearrangements with a distinct impact on protein stability. The additional residues from the VDAIKEK splicing increase the trimerisation surface which in turn bolsters the stability of the clathrin trimer. The SAS alternative splicing replaces the disordered segment with a short α-helical extension, but this apparently does not impact trimeric protein stability.

### Short alternative splicing regulates the binding of SNX5 to CHC17

In addition to mediating trimerisation, the clathrin TxD plays a key role in binding clathrin protein partners. This includes the formation of helices involved in CLC association with CHC17 and interaction with SNX5 for CHC22, the latter being essential for CHC22 subcellular localisation^6,28^. The impact of CHC17 alternative splicing on such protein-protein interactions was thus investigated through *in vitro* binding assays.

Binding to CLC was first assessed by immobilising human CHC17-FL and murine CHC17-SAS Hub fragments on Ni-NTA beads to test for capture of recombinant CLC. For this experiment, the CLCb isoform that is expressed in non-neuronal tissue was used as prey as a representative example of a CLC, given that the CHC-binding region of all CLC isoforms is functionally conserved^29^ (Fig. 5A). Both Hubs were found to bind CLCb and comparing the binding across a range of molar ratios indicates that they share similar affinities for CLC, implying that CHC17 alternative splicing does not directly modulate CLC binding.

**Figure 5.**
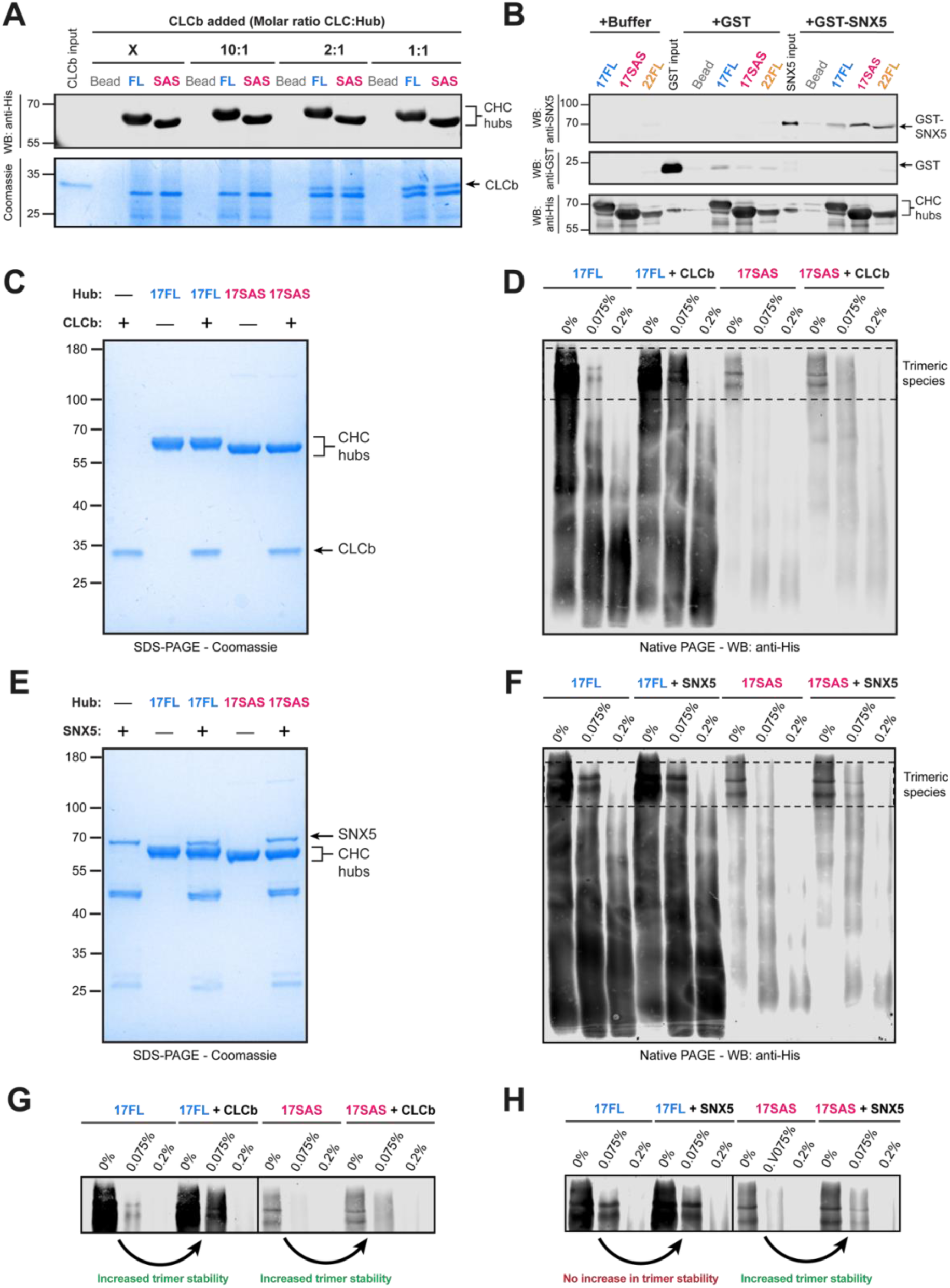
CHC17-SAS binds to SNX5 increasing trimerisation stability. **(A)** In vitro pulldown of CLCb binding to Ni-NTA bead only (Bead), CHC17-FL (FL) and CHC17-SAS (SAS). No CLCb (X) or CLCb were added at 10:1, 2:1 and 1:1 molar ratios (CLC:Hub). Samples were immunoblotted for His-tag (top), or Coomassie stained for CLCb (bottom). Position of molecular weight markers indicated on the left **(B)** In vitro pulldown of Buffer, GST or GST-SNX5 binding to Ni-NTA bead only (Bead), CHC17-FL (17FL), CHC17-SAS (17SAS) or CHC22-FL (22FL). Samples were immunoblotted for SNX5 (top), GST (middle) or His (bottom). Position of molecular weight markers indicated on the left. **(C)** Coomassie-stained SDS PAGE gel of CHC17-FL Hub (17FL), CHC17-SAS Hub (17SAS) with or without CLCb. Hub constructs comprise residues 1074-1675 for FL or 1074-1646 for SAS. Position of molecular weight markers indicated on the left. **(D)** Native PAGE gel of CHC17-FL Hub (17FL), CHC17-SAS Hub (17SAS) with or without CLCb, incubated with 0%, 0.075% or 0.2% SARC. Samples were immunoblotted for His tag. Dashed box region indicates the region of the trimeric species. **(E)** Coomassie-stained SDS-PAGE gel of CHC17-FL Hub (17FL), CHC17-SAS Hub (17SAS) incubated with or without GST-SNX5. Hub constructs include residues 1074-1675 for FL or 1074-1646 for SAS. Position of molecular weight markers indicated on the left. **(F)** Native PAGE gel of CHC17-FL Hub (17FL), CHC17-SAS Hub (17SAS) with or without GST-SNX5, incubated with 0%, 0.075% or 0.2% SARC. Samples were immunoblotted for His tag. Dashed box region indicates the region of the trimeric species. **(G) /(H)** Insets from immunoblotting in **(D)** and **(F)** to show the trimeric zone

Similarly, binding to SNX5 was examined by immobilising CHC17-FL, CHC17-SAS and CHC22 Hubs as bait to pulldown GST or GST-SNX5 (Fig. 5B). As expected, SNX5 bound to CHC22 Hub but not CHC17-FL Hub^6^. However, unlike the CHC17-FL, CHC17-SAS Hub bound SNX5. This shows that the alternative splicing at the TxD affects the protein-protein interactions of clathrin, enabling SNX5 as a new binding partner for the CHC17-SAS variant and conferring CHC22-like properties, providing a potential molecular link to GLUT4 traffic at ERGIC membranes.

CLCs play a multi-faceted role in clathrin function, including stabilising CHC17 trimerisation^18^. To explore whether the binding of SNX5 acts in the same way, the trimeric stability of CHC17-FL and CHC17-SAS Hubs was assessed in the presence or absence of SNX5 or CLCb (Fig. 5C-H). Immunoblotting shows that there is increased intensity of the trimeric species at 0.075% SARC in the presence of CLCb compared to when it is absent for both CHC17-FL and CHC17-SAS Hubs (Fig. 5D, G), confirming previously published observations that CLCs increase CHC17 trimer stability. Notably, comparing the trimer species intensity at 0.075% SARC with or without SNX5 shows that SNX5 stabilises the CHC17-SAS Hub trimer, while exhibiting no effect on CHC17-FL Hub (Fig. 5F, H). This data supports the binding studies (Fig. 5A, B) showing that the short alternative splicing does little to directly impact CLC binding, but enables the binding of SNX5 to CHC17-SAS, which in turn, increases trimer stability.

### High-resolution structure of the CHC22 TxD resembles that of CHC17 with differently charged binding surfaces

To elucidate the molecular basis for the differential binding of SNX5 to CHC TxDs, the structure of CHC22 TxD was solved to a resolution of 2.3 Å using X-ray crystallography (Fig. 6, Supplementary Table 3). CHC22, like CHC17-SAS, is truncated at the C-terminus relative to CHC17-FL. Consistent with AlphaFold predictions (Fig. 4B), the CHC22 TxD forms a predominantly α-helical trimer characterised by an extended C-terminal “tripod” helix that protrudes below the core complex (Fig. 6A). The trimer exhibits pseudo three-fold symmetry and superposition of the three monomers reveals minimal conformational variance (maximum RMSD 0.51 Å over 104 Cα atoms) (Fig. 6B). Furthermore, aligning the CHC22 TxD with the highest-resolution CHC17 TxD structure currently available (PDB: 6SCT) confirms that the overall architectural fold is conserved (Fig. 6C, D) – as evidenced by a TMscore^30^ of 0.94 for residues 1546 – 1623. The most pronounced deviation from the CHC17-FL structure is localised at the N-terminus of the CHC22 TxD where the first helix is folded back on itself to form additional contacts (Fig. 6D). This results in the TMscore dropping to 0.83 for residues 1520 – 1623. This folding is unexpected as the proximal leg should extend outwards from the trimerisation domain to form the characteristic triskelion structure and likely arises from crystal packing forces. Ultimately, the CHC22 TxD crystal structure and its comparison with CHC17-FL TxD reveal no obvious signs of a large-scale conformational rearrangement that would explain the differential binding of SNX5.

**Figure 6.**
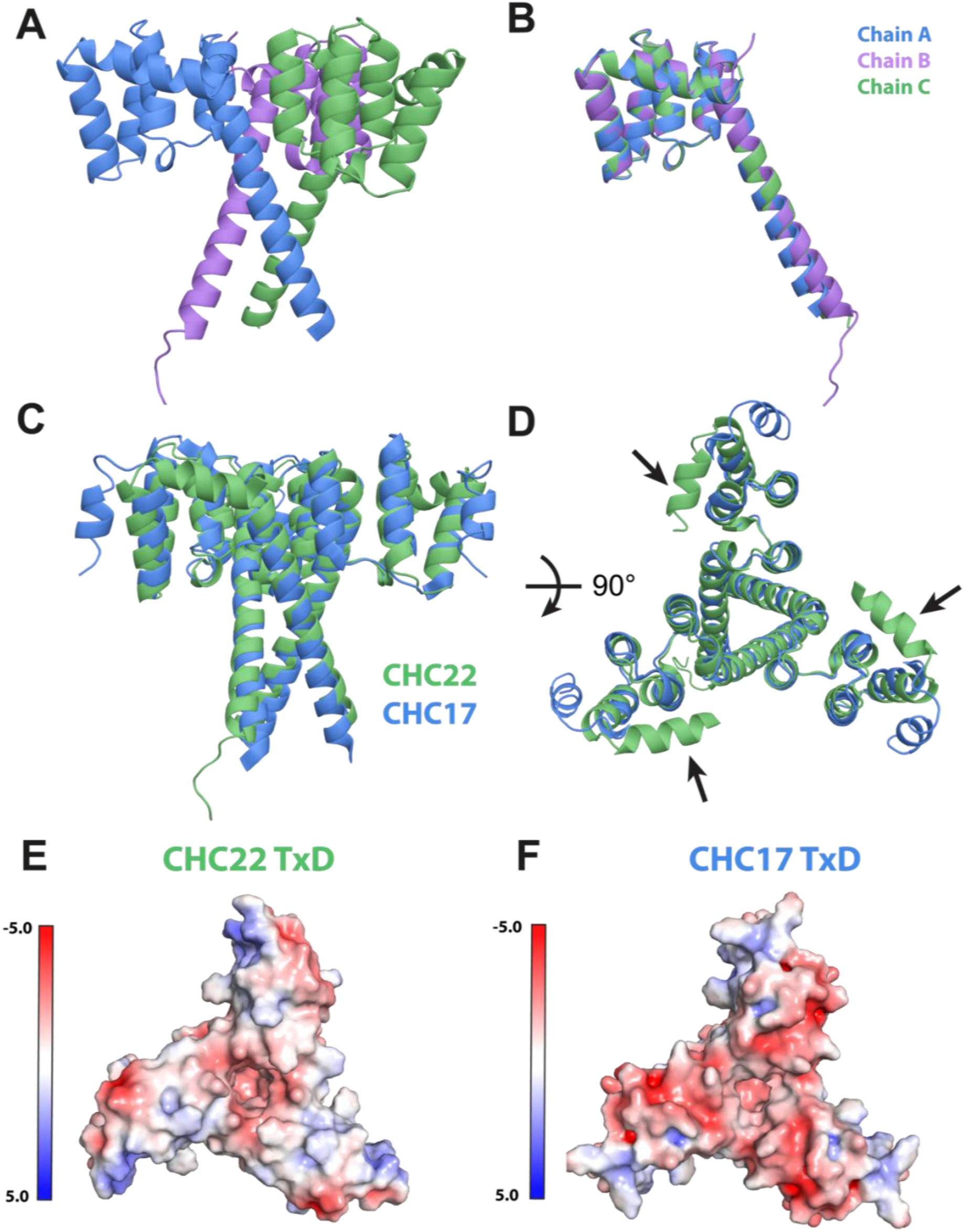
Crystal structure of CHC22 TxD highlights conserved clathrin trimerisation and distinct surface charge properties relative to CHC17. **(A)** Crystal structure of CHC22 TxD solved to a resolution of 2.3 Å, colored by chain in blue, purple or green. CHC22 TxD construct comprises residues 1520-1640. PDB Accession code: 32SP. **(B)** Overlay of each of the 3 individual chains comprising CHC22 TxD. **(C)** Structural alignment of CHC22 TxD crystal structure solved here (green) with the equivalent region of CHC17 TxD (blue) taken from a CryoEM structure of a CHC17 basket (PDB: 6SCT)^27^. Alignment calculated using MM align. **(D)** A 90° rotated view of the structural alignment in **(C)**. Arrowheads indicate the folded back N-terminal helices. **(E) /(F)** Electrostatic charge profiles of the CHC22 and CHC17 TxDs viewed from the top view shared in **(D)**. Folded back helices highlighted in **(D)** have been removed and both constructs comprise residues 1531-1626. Electrostatics calculated using APBS electrostatics with PDB2PQR and colored from −5.0 kT/e (red, negative) to +5.0 kT/e (blue, positive).

Calculating the surface electrostatics shows that the charge profiles of CHC17 and CHC22 are strikingly different (Fig. 6E, F). Previous AlphaFold modelling suggests that the ‘top’ surface of the TxD is thought to mediate interaction with SNX5^6^. This surface on CHC22 is generally neutral in its charge, with only a few minor positively and negatively charged hotspots. In contrast, the equivalent region of CHC17 is highly negatively charged – a property which would clash with the negative charged nature of the predicted interface on SNX5 (Supplementary Fig. 2A, B). This would provide an explanation for the specificity of SNX5 binding preferentially to CHC22 over CHC17-FL, but cannot immediately explain why CHC17-SAS (which shares the CHC17 FL surface sequence) binds SNX5. Notably, His-tag immunoblotting from native gels shows that the His-tag is less accessible, with consistently decreased signal for CHC17-SAS compared to CHC17-FL (Fig. 5D, F) despite SDS-PAGE gels demonstrating that the protein levels are equivalent (Fig. 5C, E). This hints that the truncation of CHC17 at its C-terminus may cause a conformational rearrangement of the N-terminus where the His-tag is located. It is possible that the C-terminal truncation of CHC17 SAS serves as a structural switch that modulates SNX5 binding, potentially by inducing a conformational change that alters the binding interface through a shift in surface electrostatics.

### CHC17 SAS compensates for the loss of CHC22 in insulin-stimulated GLUT4 traffic

Both the genetic and biochemical analysis of CHC17-SAS suggests that it may function as an alternative for CHC22. To test this hypothesis, the function of CHC17-SAS in insulin-stimulated GLUT4 membrane trafficking was investigated. In previous studies, it has been demonstrated that HeLa cells express CHC22 (anomalously for their tissue origin) and that when transfected to express GLUT4, it is packaged into a functional GSC that releases GLUT4 to the cell surface in response to insulin^4,22,31^. To test CHC17-SAS function, HeLa cells were transiently transfected with GLUT4 tagged with mCherry at the N-terminus and with an HA tag inserted in its exofacial loop, a construct that has been verified to behave as unmodified GLUT4^4^. When exported to the cell surface, the HA tag becomes accessible to anti-HA antibody detection, as a marker of translocation, compensating for the lack of antibodies recognizing the extracellular domain of GLUT4. Along with the tagged GLUT4, cells were co-transfected with siRNA targeting CHC22 (or scrambled siRNA) and one day later co-transfected with GFP-tagged CHC22, murine CHC17-FL, murine-CHC17-SAS or no CHC construct. After 48 hours, cells were treated with insulin and GLUT4 translocation was measured by binding of anti-HA antibody, prior to cell fixation to visualise the transfected CHC constructs (Fig. 7A-E). Visual analysis of cells expressing CHC constructs showed that relative to translocation in cells treated with scrambled siRNA, those treated with siRNA targeting CHC22 were impaired in GLUT4 translocation following insulin treatment (Fig. 7A, B). Translocation was rescued in cells co-transfected with GFP-CHC22 and murine GFP-CHC17-SAS (Fig. 7C, E), but not in cells co-transfected with murine CHC17-FL (Fig. 7D). Analyzing HA surface staining coincident with GLUT4 fluorescence (Manders coefficient), cells from three separate experiments were quantified. To compare between experiments, results were assessed as fold-change with insulin treatment compared to overlap under basal conditions (Fig. 7F). This quantification validated the visual analysis, showing a clear trend that insulin stimulated GLUT4 translocation is impaired with the loss of CHC22 and can only be rescued by the re-introduction of CHC22, and CHC17 SAS, but not CHC17 FL.

**Figure 7.**
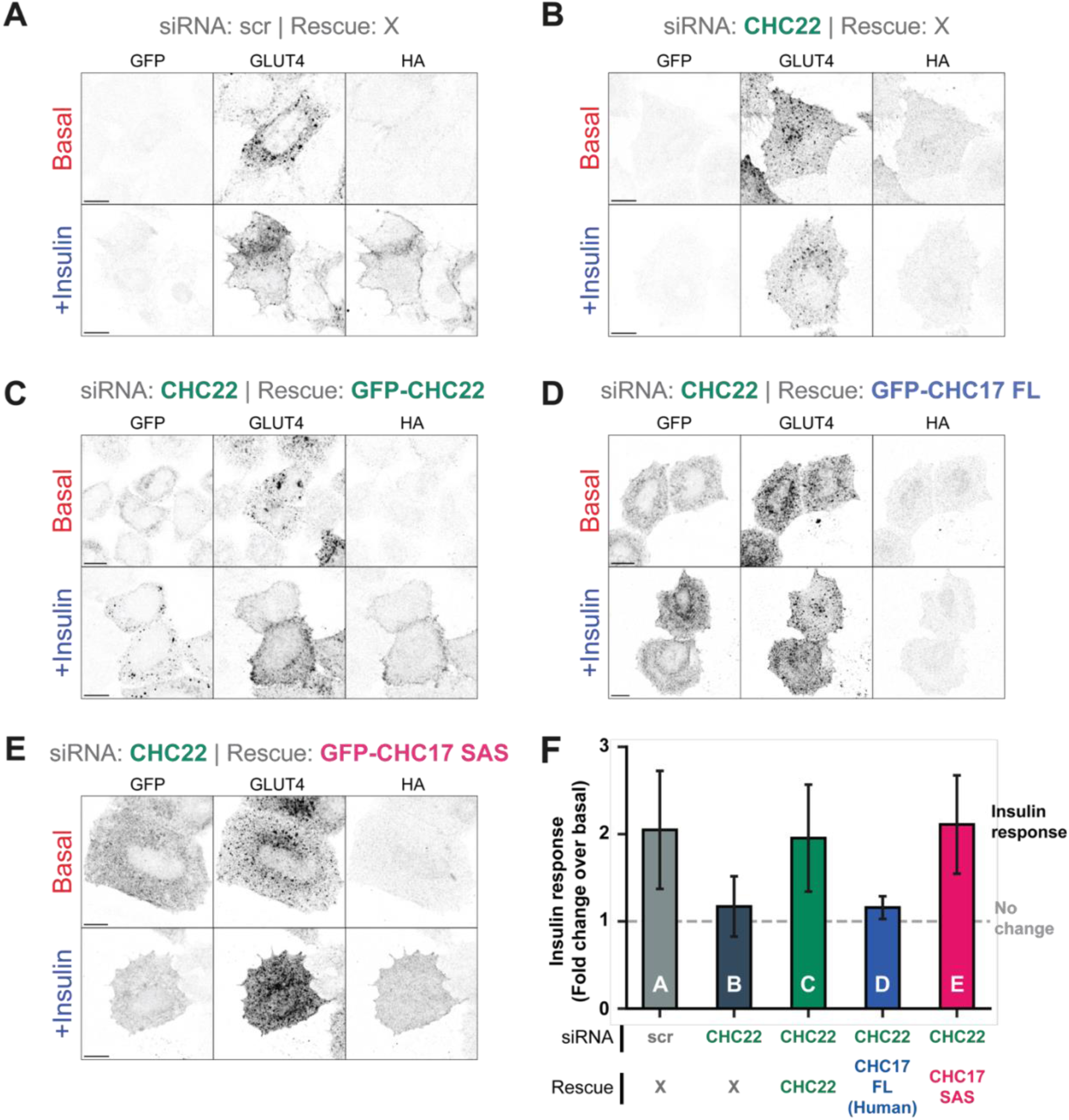
CHC17-SAS, but not CHC17-FL, rescues insulin-stimulated GLUT4 translocation following CHC22 depletion using immunofluorescence assay. Immunofluorescence-based GLUT4 translocation assay. In control conditions, HeLa WT cells were transfected with non-specific siRNA (scr) **(A)** or were depleted of CHC22 via siRNA and mock-transfected **(B)** or transfected with GFP-CHC22 (siRNA-resistant) **(C)**, GFP-CHC17-FL (murine) **(D)** or GFP-CHC17-SAS **(E)** constructs. HA-GLUT4-mCherry was additionally transfected to all cells to measure the rescue of insulin-stimulated trafficking of GLUT4 to the cell surface. Cells were live stained with anti-HA antibody targeting the surface-accessible extracellular loop of GLUT4 either before (basal) or after insulin treatment. **(A)** – **(E)** Representative immunofluorescence images of transfected and treated cells showing GFP (CHC rescue constructs), GLUT4 and HA. **(F)** The magnitude of the insulin-stimulated GLUT4 translocation as the fold-increase of colocalization between HA and GLUT4 in the insulin treated condition over the basal condition as quantified by Manders’ coefficient (n=3). Bars labelled according to conditions in **(A)** – **(E)**, error bars represent standard error of the mean.

Given the limited number of cells that can be analyzed manually, a second model GLUT4 transport system was used to further test whether CHC17-SAS can substitute functionally for CHC22 in a population of cells. HeLa cells stably transfected with HA-GLUT4-GFP were depleted of endogenous CHC22 using siRNA and insulin-stimulated translocation of GLUT4 to the plasma membrane was measured using flow cytometry to detect cell-surface anti-HA staining. siRNA-treated cells were transfected to express CHC22, murine CHC17-SAS, murine CHC17-FL and human CHC17-FL (Fig. 8). In this experiment, rescue constructs were tagged with mApple instead of GFP. Consistent with previous reports^4,6^, surface levels of GLUT4 increase by ∼60% upon insulin treatment. This insulin-stimulated increase was abrogated upon CHC22 depletion and rescued by mApple-CHC22 but not by transfection of mApple-CHC17-FL from either human or mouse. As in the smaller scale rescue experiments, transfection with mApple-CHC17-SAS was sufficient to restore insulin-stimulated translocation demonstrating that CHC17-SAS can functionally substitute for CHC22.

**Figure 8.**
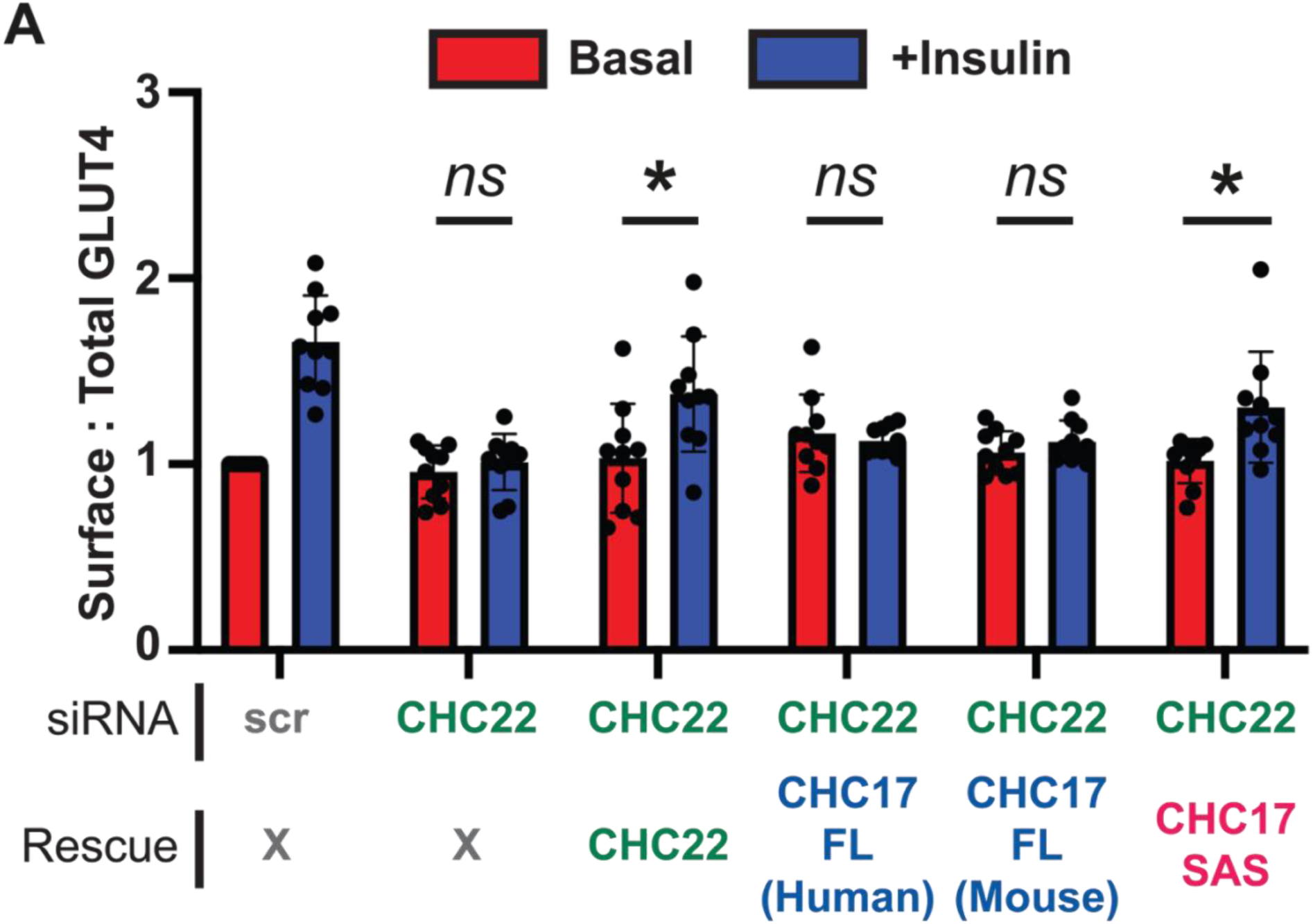
CHC17-SAS, but not CHC17-FL, rescues insulin-stimulated GLUT4 translocation following CHC22 depletion using quantitative flow cytometry assay. **(A)** Flow cytometry-based GLUT4 translocation assay. Quantification of the HA-GLUT4-GFP surface-to-total ratio (HA:GFP) using median fluorescent signal (MFI). HeLa GLUT4-GFP cells were either mock-transfected (scr) or transfected with siRNA targeting CHC22 plus constructs encoding siRNA resistant mApple-CHC22 (CHC22), Human mApple-CHC17 (CHC17-FL Human), Mouse mApple-CHC17 (CHC17-FL Mouse), and Mouse alternatively spliced mApple-CHC17-SAS (CHC17-SAS). These were then treated with vehicle only (Basal) or insulin (+Ins). Data for each experiment was normalized so that scr Basal condition equal 1. Bar height represents the mean value for all experiments in each condition. Each dot represents the normalized MFI value, for all cells in each individual experiment. Statistical analysis was performed using a one-way ANOVA with Dunnett post-hoc test (n = 10, 50,000 cells per condition, per experiment). Error bars represent standard error of the mean. ns = not significant, *P < 0.05.

## Discussion

This study aimed to investigate the mechanistic differences of GLUT4 membrane traffic across mammalian evolution, and in particular, understand how rodents target GLUT4 to the GSC in the absence of CHC22 clathrin, the protein that carries out this function in humans. Prior genetic analyses had shown that *CLTCL1* (encoding CHC22), which emerged by gene duplication at the onset of vertebrate evolution, has been lost several times throughout Mammalia, including in mice, rats and some ruminants^9,10^. Advances in sequencing technology have broadened the pool of high-quality mammalian genomes and transcriptomes available for secondary analyses. Here, the presence of *CLTCL1* was assessed across an expanded dataset of 121 mammalian genomes to find that *CLTCL1* was lost in at least 4 mammalian lineages. This validates previous genetic analyses and further demonstrates that the loss of *CLTCL1* is a prevalent, independently occurring feature of the Rodentia, Cetartiodactyla, Chiroptera and Monotremata orders.

Transcriptomic analysis showed that species that have lost *CLTCL1* instead express an alternatively spliced variant of *CLTC*, which is truncated at the extreme C-terminus, mirroring the truncation of *CLTCL1*. The alternatively spliced variant (*CLTC-SAS*) displays a similar tissue-specific distribution (predominantly in muscle) and the transcript is present at similar levels to *CLTCL1*, relative to *CLTC*. This mutually exclusive expression is striking, with all investigated species expressing *CLTCL1* or *CLTC-SAS*, but never both. This phenomenon of an ancestral protein retaining a broad expression pattern, while the duplicate is restricted to specific cell types is a common evolutionary fate for duplications and alternatively spliced isoforms^32^.

We further demonstrate here that the protein resulting from the murine *CLTC-SAS* transcript (CHC17-SAS) is expressed in mouse muscle tissue (and not kidney) at levels relative to CHC17-FL comparable to the CHC22/CHC17 ratio in human muscle, confirming tissue specificity of the identified splice variant, mirroring the trend observed at a transcript level. The newly characterised alternatively spliced isoform CHC17-SAS alters the CHC17 protein within the clathrin trimerisation domain (TxD), the same domain that is altered in another alternatively spliced version of CHC17 (CHC17-VDAIKEK) in murine and human muscle^17^. Protein modelling and biophysical characterisation showed that the additional residues spliced into CHC17-VDAIKEK improve clathrin trimer stability, while the truncation characteristic of CHC17-SAS has little effect. However, analysis of protein-protein interactions found that the TxDs of two ‘truncated’ CHCs (CHC22 and murine CHC17-SAS) bind to SNX5 in contrast to the elongated CHC17-FL, which does not interact with SNX5. In humans, SNX5 provides a bridge between the CHC22 and p115 and is required for CHC22 recruitment to the ERGIC, suggesting that the ability of CHC17-SAS to bind SNX5 could enable it to perform a function equivalent to that of CHC22.

Previously published protein modelling suggested that SNX5 binds to the exposed ‘top’ surface of CHC22, perpendicular to the helix tripod that CHC17 utilises to interact with its regulatory CLC subunit^6^. To further understand this interaction, the structure of CHC22 TxD was solved using X-ray crystallography. This experimentally determined structure demonstrated that CHC22 and CHC17 have differing electrostatic profiles on the ‘top’ surface that could explain the molecular specificity of SNX5 for CHC22 over CHC17. However, it remains unclear how CHC17-SAS, which has the same sequence as CHC17-FL is able to bind to SNX5 while CHC17-FL cannot. It is possible that the truncation in CHC17-SAS shifts the position of the tripod helices to alter the conformation of this ‘top’ surface and expose SNX5 binding residues. Such alterations are not visible using AlphaFold to model CHC17-SAS, as modelling is based on existing structures. Alternatively, the disordered C-terminal extension, unique to CHC17 FL, could reach up to the top surface of CHC17 to prevent SNX5 from binding. Such autoinhibition by disordered regions have been observed in many other contexts^33^. Finally, we cannot rule out the possibility that the binding of SNX5 to CHC22 is not as predicted by AlphaFold and is instead at this ‘bottom’ surface at the extreme C-terminus, where the truncation relative to CHC17 in CHC22 and CHC17-SAS occurs. Nonetheless, the protein interaction data indicates that CHC17-SAS can bind both CLC and SNX5, and the latter binding would enable recruitment of CHC17-SAS to the intracellular site where CHC22 functions in humans. These data further support a role for the clathrin TxD in influencing clathrin flexibility. Although the Hub regions of CHC17 appear unchanged in cryo-EM models of differently-sized clathrin baskets^27^, the inclusion of the VDAIKEK sequence by splicing was shown to promote flat clathrin lattice formation^17^. Thus, tweaking the TxD may indeed influence clathrin structure.

Previous studies showed that engineering mice to express exogenously introduced *CLTCL1* in addition to their endogenous *CLTC-SAS* in muscle resulted in a situation where GLUT4 was over-sequestered and trapped inside an enlarged GSC and unable to be mobilised to the cell surface^5^. These findings already suggested functional redundancy of murine and human GLUT4 trafficking pathways. Consistent with its ability to bind SNX5, we demonstrate here that murine CHC17-SAS, but not human or murine CHC17-FL, can substitute for CHC22 in two independent HeLa cell models of insulin-stimulated GLUT4 trafficking. When CHC22 was depleted by siRNA in both models, insulin-responsive GLUT4 translocation was lost, then regained by expression of either CHC22 or CHC17-SAS. Together these findings allow us to propose a common model for GLUT4 trafficking in Mammalia where newly synthesised GLUT4 is trafficked to the GSC from the ERGIC by a truncated CHC represented by CHC22, where it is present or CHC17-SAS, when CHC22 is absent.

An outstanding question remains as to the evolutionary drivers of utilising two distinct genes compared to using one single gene that is alternatively spliced and how this transition occurred across multiple vertebrate lineages. Instances of isoform switching as a compensatory mechanism in response to paralogue loss have been characterised at single cell resolution^34^, and in well-defined systems such as the Muscleblind-Like (MBNL) family of proteins^35^. In contrast, other evolutionary analyses suggest that in many cases following gene duplication, the newer duplicates may take over protein functions that were previously controlled by alternative splicing^36^. However, interspecies examples of specific compensatory mechanisms have not been broadly discussed in previous literature.

One potential benefit of utilising two distinct genes is that it allows for greater specialisation of clathrin isoforms. In humans, CHC17 and CHC22 share 85% sequence identity but have specific points of divergence that allow their deviation into distinct functions. In fact, this is often a selective pressure that results from gene duplication to avoid interactions with the duplicated partner^37^. For example, CHC22 has a patch within its N-terminal domain that is divergent from CHC17 that allows it to directly interact with p115 for ERGIC recruitment, while CHC17 cannot. Furthermore, CHC22 does not bind CLCs in a cellular context. This enhances the specificity of membrane recruitment, facilitating the two-part mechanism exhibited by CHC22, requiring both direct interactions with p115 via the CHC22 N-terminal domain as well as the indirect interaction with p115 via SNX5^6^. As a result, CHC17 and CHC22 are rarely found within the same clathrin coats^8^ and their respective membrane trafficking functions can be regulated independently. In contrast, where CHC17 is alternatively spliced, CHC17-FL and CHC17-SAS have identical sequences up until the final residues, making differentiation more challenging. CHC17-SAS does not have the patch in the terminal domain to additionally bind p115 and its ability to bind CLCs may affect SNX5 binding. This adds up to the suggestion that the role for CHC17-SAS in direct GLUT4 targeting to the GSC may be reduced in mice relative to its role in humans, though p115 is implicated in GLUT4 targeting in mice^12^. Consequently, the endocytic retrograde route for GLUT4 return to the GSC is predicted to play a dominant role in formation of the murine GSC, which would function through AP1 binding by CHC17-SAS, similarly to the CHC22-AP1 pathway. It is possible that species that maintain the *CLTCL1* gene require more stringent regulation of their glucose metabolism and require the enhanced regulation that possessing two genes provides, while species that lack *CLTCL1* favour the simplicity of a single gene and compensate for some functional cross-over by emphasising different arms of the GLUT4 trafficking pathway.

If the evolutionary pressure to lose *CLTCL1* in favour of *CLTC-SAS* relates to efficiency, then it is important to establish why this is apparently so strong in the Rodent and Cetartiodactyla lineages in particular, which appear to have evolved *CLTC-SAS* independently as an example of convergent evolution using alternative splicing. One clue may lie in the conservation of *CLTCL1* in the bats of the Chiroptera order, split between the old-world bats (*CLTCL1* present) and new-world bats (*CLTCL1* absent). Investigating what evolutionary pressures or dietary restrictions these groups have in common may hint at what independently drove the adoption of *CLTC-SAS* over *CLTCL1* in these groups in particular.

This work elucidates the mechanism of GLUT4 trafficking in mammalian species, including mouse (*M. musculus*) and rat (*R. norvegicus*). These species are of particular relevance due to their prevalent use as preclinical models for metabolic diseases such as Type 2 diabetes (T2D). CHC22 is a potential drug target for T2D, found to be highly enriched in the GSCs of T2D patients, potentially exacerbating symptoms by over-sequestering GLUT4 inside the cell^5^. However, it was not previously clear how this disease phenotype related to mouse models, which do not have CHC22. By identifying CHC17-SAS as a surrogate for CHC22 in mice, this points to a common mechanism of GLUT4 trafficking, opening avenues for shared features of CHC22 and CHC17-SAS to be targeted and clinically studied in human and murine models. For example, both CHC22 and CHC17-SAS lack the ‘QLMLT’ motif within the trimerisation domain that is required for CHC17 clathrin uncoating. Uncovering the potentially shared mechanism by which these truncated CHCs are uncoated could form the basis for therapeutic treatments against T2D through the stimulation of uncoating. This would reverse the over-sequestration of GLUT4 in diabetes patients, removing CHC22 from the GSC to allow more GLUT4 to mobilise to the cell surface.

## Materials and Methods

### Genomic datasets and analysis

Mammalian genome sequences were downloaded from Ensembl genome browser (https://www.ensembl.org/index.html, accessed: January 2022) and filtered to include only those fitting each of the following criteria: >30x average coverage, >99% ungapped length, N50 >100000, L50 <1000 and total scaffold count <20,000. To determine presence and location of *CLTC* and *CLTCL1* within the 121 species from this filtering, homologous sequence corresponding to human CHC17 and CHC22 was searched by tBLASTn^38^ (Supplementary Table 1). Corresponding sequence was extracted and the unrooted phylogenetic tree was generated with MEGA 11^39^ using extracted *CLTC* sequences.

### Transcriptomic datasets and analysis

Paired-end RNA-seq datasets representing 18 different mammalian species and nine tissue types were identified and downloaded from Sequence Read Archive (SRA) – listed in Supplementary Table 4. Sequences were trimmed using Trimmomatic (v0.39)^40^ and aligned against contemporaneous genome assemblies and annotations using hisat2 (v2.2.1)^41^. The location of final exon splicing junctions, in addition to a ‘control’ junction shared by all major *CLTC* splicing isoforms, were identified manually within each genome assembly. Exon junction count analysis utilised the featureCounts facility in the ‘Rsubread’ R package^42^. For rat and mouse datasets, where two alternative exon junctions exist in the alternatively spliced transcript, the maximum exon junction count was utilised.

### Cell culture and transfection

Cell lines were cultured at 37 °C in a 5% CO2 atmosphere with 100% humidity. Cells were grown in DMEM high glucose (Gibco) supplemented with 10% FBS (Gibco), 50 U/ml penicillin and 50 µg/ml streptomycin (Gibco). The HeLa lines used in this study were as follows: HeLa Wild-type (WT) for colocalization and HeLa HA-GLUT4-GFP (HeLa-G4) for FACS^4,6,22,31^. Both cell lines used tested negative for mycoplasma infection.

For transfection of siRNA, oligonucleotides were complexed with jetPRIME (Polyplus) in accordance with manufacturer’s instructions and added to the cells to a final concentration of 20 nM for all targets. For transfection of DNA plasmids, cells were seeded at the same density and the following day, were transfected with plasmid DNA which was complexed with jetPRIME in a 1:2 mixture (DNA/jetPRIME) in accordance with manufacturer’s instructions.

For both transfection types, cells were then returned to normal growth conditions and processed downstream as described.

### siRNA

siRNA targeting CHC22 or nontargeting control siRNA as published previously^4,6^.

### Plasmid constructs

For GLUT4 translocation assays with immunofluorescence or flow cytometry, plasmids encoding HA-GLUT4-mCherry, GFP-CHC and mApple-CHC constructs were utilised. HA-GLUT4-mCherry and mApple-CHC22 were generated as described in previous studies^6,10^. mhCHC17 (human) was subcloned into the mApple-CHC22 backbone from eGFP-CHC17 plasmid utilised previously^7^. To generate mApple-mCHC17 (mouse) FL, Q5 site directed mutagenesis was used to convert differing residues to their mouse equivalent. To generate mCHC17 SAS, the DNA corresponding to the truncated region was deleted and replaced with DNA encoding the alternative final exon through a HFi assembly reaction (NEB). GFP-CHC constructs were generated by restriction cloning into a sfGFP pcDNA5 backbone according to manufacturer’s instructions (NEB).

### Antibodies

The primary antibodies in this study were used at a dilution of 1:200-1:500 for IF, 1:2000-1:5000 for immunoblotting or as described for immunoprecipitation. The primary antibodies used in this study include: mouse anti-CHC17 (X22) for immunoprecipitation (20 µg/reaction); mouse anti-HA (BioLegend 901503) for live-cell surface labelling of GLUT4 for immunofluorescence and flow cytometry translocation assays; rabbit anti-GLUT4 (Abcam Ab313775) for immunofluorescent labelling of total GLUT4 in GLUT4 translocation assay; chicken anti-GFP (Invitrogen A10262) for immunofluorescent labelling of GFP-tagged rescue constructs in translocation GLUT4 assay; mouse anti-His (R&D systems MAB050H) for immunoblotting analysis of recombinant protein pulldown assays and Native-PAGE assays. Secondary antibodies were used as follows: For IF, species-specific secondaries conjugated to, Alexa Fluor 488 (anti-chicken Invitrogen SA1-72000), Alexa Fluor 555 (anti-rabbit Invitrogen A21428), or Alexa Fluor 647 (anti-mouse Invitrogen A331571) were used at a dilution of 1:1000. For fluorescent western immunoblotting, secondaries conjugated to IRDye were used, anti-mouse 680RD (LI-COR Biosciences 926-68070) at a dilution of 1:10,000.

### GLUT4 translocation using immunofluorescence

Hela WT cells were seeded on coverslips in 24-well plates and transfected with CHC22 siRNA 24 h later. After a further 24 h, cells were transfected with HA-GLUT4-mCherry and rescued via transfection with GFP-CHC before incubation for a further 48 h. On the day of experimentation, cells were washed once with HBSS (Gibco) and incubated in HBSS for 10 min. Control cells (basal) were stained at this point (see below). HBSS was replaced with DMEM without serum containing 1 µM insulin (Sigma) or the same volume of vehicle (water) and then incubated for 20 min. The cells were then placed on ice and washed with PBS containing magnesium and calcium ions (PBS^+/+^) (Corning) which had been pre-cooled to 4°C, for a total of three times. The cells were then live-stained by incubating with mouse anti-HA antibody solution (1:500 diluted in DMEM without serum; 0.002 μg/mL) on ice for 30 min to detect surface GLUT4. After this, cells were washed three times in PBS^+/+^ and fixed in 4% PFA for 15 min on ice. After fixation, the cells were permeabilized in PBST (PBS with 0.1% Triton X-100) for 5 min. Coverslips were washed three times in PBS at room temperature and then blocked in PBS with 5% FBS for 1 h. Blocked coverslips were incubated with additional primary antibodies (rabbit anti-GLUT4), chicken anti-GFP diluted in PBS + 0.1% FBS overnight at 4 °C. Coverslips were washed three times in PBS then incubated with secondary antibody solution (anti-chicken Alexa Fluor 488, anti-rabbit Alexa Fluor 555 or anti-Mouse Alexa Fluor 647) diluted 1:1000 in PBS + 0.1% FBS for 1 hour at room temperature followed by washing three times in PBS. DAPI staining (1:10000 in PBS) was performed for 10 min and coverslips washed three times in PBS. Coverslips were mounted on microscope slides (Superfrost, Thermo Fisher Scientific) using 5μl of Diamond Vectashield per coverslip (Thermo Fisher Scientific) and set overnight at 4 °C.

#### Image acquisition and analysis

All samples were imaged using a Leica TCS SP8 inverted laser-scanning confocal microscope equipped with two high-sensitivity hybrid detectors and one photomultiplier. A 63x (1.40 NA) HC Plan-Apo CS2 oil-immersion objective was used with five laser lines. Fluorophores were sequentially excited at 405 nm (DAPI), 488 nm (GFP, Alexa Fluor 488), 543 nm (mApple, Alexa Fluor 555), 561 nm (Alexa Fluor 568), and 633 nm (Alexa Fluor 647). For fixed images, 8-bit images were acquired (1024×1024 XY pixel aspect). For each image, a Z-axis sectional stack of 8 sections was acquired using the GLUT4 signal as a positional marker for start and end positions. The images were saved in .tiff and exported for further processing. For visualisation, the images were processed as individual sections and analysis performed using Fiji^43^ to obtain image embedded data. For colocalization, a Python script was written for Napari integration to enable analysis in only selected cells that expressed sufficient levels of HA-GLUT4-mCherry and, in the CHC rescue conditions, also expressing enough GFP. For the cells in the region of interest (ROI) a mask was applied either by manual or automatic threshold. Threshold was selected to include all pixels with a positive signal for a channel, later subjected to a colocalization analysis for Pearson’s and Manders coefficients between two channels. All sections of each image were summed and analyzed. Cells were only included if biologically relevant distribution and quantity of GLUT4 and CHC was observed (overexpression and ER distribution excluded). The Manders coefficient (M2) was calculated as the fraction of HA signal (surface GLUT4) that overlaps with GLUT4 (total GLUT4). Details regarding code availability are provided as part of the data availability statement.

### GLUT4 translocation using flow cytometry

HeLa HA-GLUT4-GFP (HeLa-G4) cells were seeded in 6-well plates at standard density and, the following day, transfected with siRNA for KD (knockdown) and incubated for a total of 72 h thereafter. For the rescue conditions using mApple tagged CHC plasmids, the cells were re-transfected with expression plasmids in fresh media at the 48 h post KD time point and processed 24 h later, at the 72 h endpoint for all FACS experiments. On the day of the experiment (72h post KD, 24h post plasmid transfection), cells were washed (three times, PBS, 37 °C) and serum starved in EBSS (Gibco) for 2 h prior to insulin stimulation. For insulin stimulation, insulin (Sigma) or the same volume of vehicle (water) was added to the wells, to a final concentration of 170 µM, for 20 min at 37 °C. The cells were then placed on ice and washed with PBS containing magnesium and calcium ions (PBS^+/+^) (Corning) which had been pre-cooled to 4 °C, for a total of three times. The cells were then live-stained by blocking for 30 min (PBS^+/+^ + 2% horse serum) on ice and then incubating with anti-HA antibody solution (1:200 diluted in PBS^+/+^+ 2% horse serum) on ice for 1 h to detect surface GLUT4. After this, cells were washed three times in PBS^+/+^ and fixed in 4% PFA for 15 min on ice. After fixation, cells were washed three times in PBS at room temperature. Cells were then incubated with secondary antibody solution (anti-Mouse Alexa Fluor 647 diluted 1:1000 in PBS+ 2% horse serum) for 1 hour at room temperature followed by washing three times in PBS and gently lifted using a cell scraper (Corning). These suspensions were pelleted by centrifugation at 300x g for 3 min and resuspended in a 500 µL of PBS.

#### Flow cytometry data acquisition and analysis

Flow cytometry data was acquired with FACS Diva acquisition software on a LSRII flow cytometer (Becton Dickinson) equipped with violet (405 nm), blue (488 nm), yellow (561), and red (633 nm) lasers. Typically, 50,000 events were acquired, and median fluorescent intensity (MFI) values for surface GLUT4 (Alexa Fluor 647), CHC plasmid transfectants (mApple), and total GLUT4 (GFP) were recorded. Post-acquisition analysis was performed using FlowJo software (TreeStar), where debris was removed by using forward/side light scatter gating. Then fluorescence histograms were analyzed using gating to select only the transfected cell populations in the experiments based on the fluorescent signal (mApple) from the expression of the CHC fusion proteins. The MFI values were exported and processed in Excel (Microsoft), whereby the ratios of surface-to-total MFI were calculated to quantify the extent of GLUT4 translocation in each condition.

### SDS-PAGE and immunoblotting analyses

For conventional immunoblotting, protein samples were separated by SDS-PAGE (10% acrylamide, Merck Millipore), transferred to nitrocellulose membrane (0.2 µm, Bio-Rad), incubated with primary antibodies (1–5 µg/ml) overnight at 4 °C, washed, and labelled with species-specific IRDye-conjugated secondary antibodies added at a dilution of 1:10,000 (LI-COR Biosciences). These labelled membranes were imaged using the Odyssey DLx imaging system (LI-COR Biosciences). Quantification of the relative abundance of proteins in the membranes was performed using Image Studio (LI-COR Biosciences).

For Coomassie staining, SDS-PAGE gels were stained using InstantBlue Coomassie Protein Stain (Abcam) before imaging using a ChemiDoc MP imaging system (Bio-Rad).

### Immunoprecipitation of CHC17-SAS from mouse tissue

All mouse tissue used in this study were harvested from wild type (WT) animals of the C57BL/6 genetic background strain. Tissues were primarily harvested as part of another study^44^ examining tissues other than the kidney and skeletal muscle. Unused tissues from that study were used for this study, therefore no additional animals were sacrificed for the experiments undertaken in this study. Mouse tissue had been harvested and snap-frozen in liquid nitrogen and stored at −80 °C until use. All animal procedures and breeding were conducted according to the Animals Scientific Procedures Act UK (1986) and in accordance with the ethical standards of University College London.

Mouse tissue was removed from −80 °C storage and placed on ice, and lysed in cold CHC17IP lysis buffer (Supplementary Table 2) supplemented with cocktail of protease inhibitors (1 tablet/10 ml; Thermo Fisher Scientific). Tissues were lysed using Dounce homogenizer for mechanical shearing with several (up to 20 for skeletal muscle) passes of the pestle. These lysates were centrifuged (16,000 x g for 15 min, 4 °C) to remove nuclei and cellular debris. The protein content of these lysates was quantified using BCA (Pierce). The concentrations of the input lysates were adjusted and a 50 μl sample of the input lysate taken. For CHC17 immunoprecipitation from mouse tissue, 20 µg of specific anti-CHC17 antibody (X22) was incubated with 500 μl of 10-20 mg/ml of precleared post-nuclear lysates overnight at 4 °C. Samples were then incubated with washed and pre-blocked protein G Sepharose beads (50 µl, GE Healthcare) for 45 mins at 4 °C, before three consecutive washing steps in lysis buffer. The pelleted protein G Sepharose beads were then resuspended in 50 µl of 2x Laemmli sample buffer (Bio-Rad) and proceeded to SDS-PAGE.

### Mass Spectrometry

Gel pieces were digested with trypsin overnight, and the digestion was stopped adding 1% trifluoroacetic acid to a final pH of 2. Peptides were dried and dissolved in 2% formic acid before liquid chromatography–tandem mass spectrometry (MS/MS) analysis. The mixture of tryptic peptides was analyzed using an Ultimate3000 high-performance liquid chromatography system coupled online to an Eclipse mass spectrometer (Thermo Fisher Scientific). Buffer A consisted of water acidified with 0.1% formic acid, while buffer B was 80% acetonitrile and 20% water with 0.1% formic acid. The peptides were first trapped for 1 min at 30 μl/min with 100% buffer A on a trap (0.3 mm by 5 mm with PepMap C18, 5 μm, 100 Å; Thermo Fisher Scientific); after trapping, the peptides were separated by a 50-cm µPAC NEO column (Thermo Fisher Scientific). The gradient was 3 to 35% B in 19 min at 750 nl/min. Buffer B was then raised to 55% in 2 min and increased to 99% for the cleaning step. Peptides were ionized using a spray voltage of 2.1 kV and a capillary heated at 275 °C. The mass spectrometer was set to acquire full-scan MS spectra (350 to 1400 mass/charge ratio) for a maximum injection time set to Auto at a mass resolution of 120,000 and an automated gain control (AGC) target value of 100%. For a 1.2 second the most intense precursor ions were selected for MS/MS. HCD fragmentation was performed in the HCD cell, with the readout in the Orbitrap mass analyser at a resolution of 30,000 (isolation window of 1.4 Th) and an AGC target value of 200% with a maximum injection time set to Auto and a normalized collision energy of 30%. All raw files were analyzed by FragPipe v20 software using the integrated MsFragger v3.8 search engine^45^. All peptides identified were validated with Philosopher v5.0^46^ at 1% FDR and quantified with IonQuant v1.9.8^47^. All files were searched against the Human SwissProt Proteome (07/2022 release with 20,383 protein sequences). MsFragger was used with the Semi-tryptic mode with the addition of deamidation (N) as variable modification. Figures were created directly using FragPipe-PDV viewer and exported in pdf format.

### Protein expression and purification

All buffers utilised here are summarised in Supplementary Table 2.

#### CHC fragments

CHC22 TxD (residues 1520-1640) and CHC17 Hubs, including FL (residues 1074-1675), SAS (residues 1074-1646) and VDAIKEK (1074-1682) were cloned into pET15b as N-terminal 6xHis fusions with a thrombin cleavage site. CHC22 Hub (residues 1074-1640) was cloned into a vector derived from pET-Duet also as an as N-terminal 6xHis fusion with a thrombin cleavage site. CHC22 TxD and CHC17 Hubs were expressed in BL21(DE3) *E. coli* and induced with 0.25 mM IPTG at 37 °C for 4 h (Hubs) or with 0.5 mM IPTG at 30 °C for 5 h (TxD). CHC22 hub was expressed in Rosetta 2 pLysS *E. coli* and induced with 0.4 mM IPTG at 18 °C for 18 h.

For structural and biochemical studies CHC22 TxD and CHC17 Hub cell pellets were lysed by sonication in their respective binding buffers and the lysate clarified at 40,000 x g for 45 mins at 4 °C. CHC17 Hubs were affinity purified using Ni-NTA bead slurry (HisPur Ni-NTA resin; Thermo Fisher) and eluted with CHC17Hub elution buffer. CHC17Hubs were then polished via size exclusion on a Superose 6 10/300 GL column (Cytiva) in CHC17Hub SEC buffer. CHC22 TxD was affinity purified using a 1 ml/5 ml HisTrap HP column (Cytiva) and eluted using in a linear gradient from 50 mM to 400 mM imidazole (CHC22TxD elution buffer). The pooled eluate was concentrated and purified by size exclusion on a HiLoad 16/60 Superdex 75 column (Cytiva) in CHC22 TxD SEC buffer.

#### CHC interacting proteins

SNX5 (residues 1–404) was cloned as an N-terminal GST-fusion with a thrombin cleavage site in pGEX3 backbone, and induced in Rosetta 2 pLysS *E. coli* with 0.5 mM IPTG at 18 °C for 18 h. CLCb (residues 1-211) was cloned as an N-terminal 6xHis fusion with a thrombin cleavage site in a pET28 plasmid backbone and induced in BL21(DE3) *E. coli* with 1 mM IPTG at 37 °C for 4 hours.

GST-SNX5 and CLCb were purified via affinity (SNX5; GST, CLCb; Ni) and size exclusion chromatography as previously described^6,16^. Where required, the His-tag was removed from CLCb by thrombin cleavage and un-tagged CLCb purified via reverse Ni affinity purification.

### In vitro binding assays

Pulldowns with GST-only or GST-SNX5 was prey and CHC22/CHC17Hub (FL/SAS) as bait were carried out as previously described^6^. For pulldowns with CLCb (untagged) as prey and CHC17Hubs (FL/SAS) as bait, CHC17Hubs were affinity purified onto Ni-NTA beads from crude *E. coli* lysate in CHC17Hub binding buffer. For each pulldown, 20 μl of 50% bead slurry was pre-blocked in CLC PD buffer for 1 hour and incubated with CLC PD buffer only or CLCb at a 10:1, 2:1 or 1:1 molar ratio (CHC17Hub : CLCb) for 45 minutes. Beads were washed x 3 in CLC PD wash buffer and eluted into SDS-PAGE sample buffer. Pulldowns were analyzed by Western blotting using an anti-His-tag antibody and Coomassie-stained SDS-PAGE.

### Detergent challenge assay to measure Hub trimeric stability

The impact of alternative splicing and binding partners on clathrin trimer stability was assessed using a challenge assay using the detergent Sarkosyl (N-lauroylsarcosine) as previously described^18^. Purified CHC17 hub constructs (CHC17 FL, SAS, VDAIKEK) alone or complexed with CLCb or GST-SNX5 at a 1:1.1 molar ratio (Hub : partner) for 1 hour were then incubated with increasing concentrations of SARC (0.0% - 0.2% final) for 15 mins on ice. Following SARC treatment, samples were analyzed by Native-PAGE (10% resolving, 4% stacking), run at 100V for 4 hours, followed by Coomassie staining or anti-His immunoblotting

### CHC22 TxD crystal structure

CHC22 TxD crystals were obtained under two conditions (JCSG D2 and JCSG B10; JCSG+ screen, Molecular Dimensions). For the JCSG D2 condition, 150 nl CHC22 TxD (CHC22 TxD SEC buffer) at 17.5 mg/ml was mixed with 150 nl precipitant solution (JCSG D2; 0.1 M HEPES pH 7.5, 0.2 M MgCl_2_, 30% PEG-400) in a sitting-drop vapour diffusion experiment equilibrated above 75 μl precipitant solution. For the JCSG B10 condition, 175 nl CHC22 TxD in SEC buffer at 15.0 mg/ml was mixed with 175 nl precipitant solution (0.1 M sodium cacodylate pH 6.5, 0.2 M MgCl_2_, 50% PEG-200) in a sitting drop equilibrated against 75 μl of precipitant solution.

Diffraction data for both crystals were collected on beamline ID30A-1 at the European Synchrotron Radiation Facility (ESRF, Grenoble, France). Initial phases were obtained using data from the JCSG B10 crystal by molecular replacement protocol using MrBUMP^48^ with an AlphaFold prediction of the CHC17 TxD domain (AF-E9QBV1-F1), as the search model. As the crystal obtained under the JCSG D2 condition yielded higher-quality diffraction data, subsequent structure determination and refinement were performed using this dataset. The final structure reported here was solved using the model derived from the JCSG B10 dataset. The JCSG D2 crystal diffracted to 2.18 Å; however based on CC_1/2_ criterion, the final resolution cutoff was set at 2.30 Å. Model building and refinement were performed using COOT^49^, REFMAC5^50^ and PHENIX.refine^51^

### Protein modelling and calculations of trimeric binding energy and electrostatics

Models of CHC17 FL, SAS and VDAIKEK TxDs (residues 1522-end) were generated using AlphaFold-Multimer^23,24^. To predict the binding energies of the trimerisation interface, the AlphaFold generated models of CHC17 FL, SAS and VDAIKEK TxDs were first refined using the *FastRelax* protocol with thorough cycle sampling (*-relax:thorough*) in Rosetta (v3.13)^52^, generating 3 independently relaxed structures. For each model, the trimer binding energies were calculated pairwise across all three chain interfaces (AB, AC, BC for chains ABC) using Rosetta *InterfaceAnalyzer*^53^. Resulting ΔG values were averaged across the three interfaces per relaxed structure, and this value was averaged across the three independently relaxed structures to evaluate the representative trimeric binding energy per construct expressed in Rosetta Energy Units (REU).

The electrostatic potential of CHC17 and CHC22 TxDs were calculated using the Adaptive Poisson-Boltzmann Solver (APBS) plugin in PyMOL^54^. Structure files were prepared with PDB2PQR^55^ at pH 7.4 using the PARSE force field. APBS calculations were carried out at 310K and 150mM NaCl. Surface potentials were expressed between −5.0 kT/e and +5.0 kT/e.

Structural alignments and TM-score calculations between CHC22 TxD and CHC17 TxD (from PDB:6SCT)^27^ were assessed using MMalign^30,56^.

#### Statistical analysis

Statistical analysis was performed in GraphPad Prism. First, the data were cleaned using ROUT detection of outliers in Prism, followed by testing for normal distribution (D’Agostino - Pearson normality test). Then, the significance for parametric data was tested by either one-way ANOVA followed by Tukey’s or Dunnett’s post-hoc multiple comparison test, or an unpaired two-tailed t-test with Welch’s correction. Detailed statistical information for each experiment, including statistical test, number of independent experiments, p values, and definition of error bars are listed in individual figure legends.

## Summary of supplementary material

**Supplementary Figure 1.**
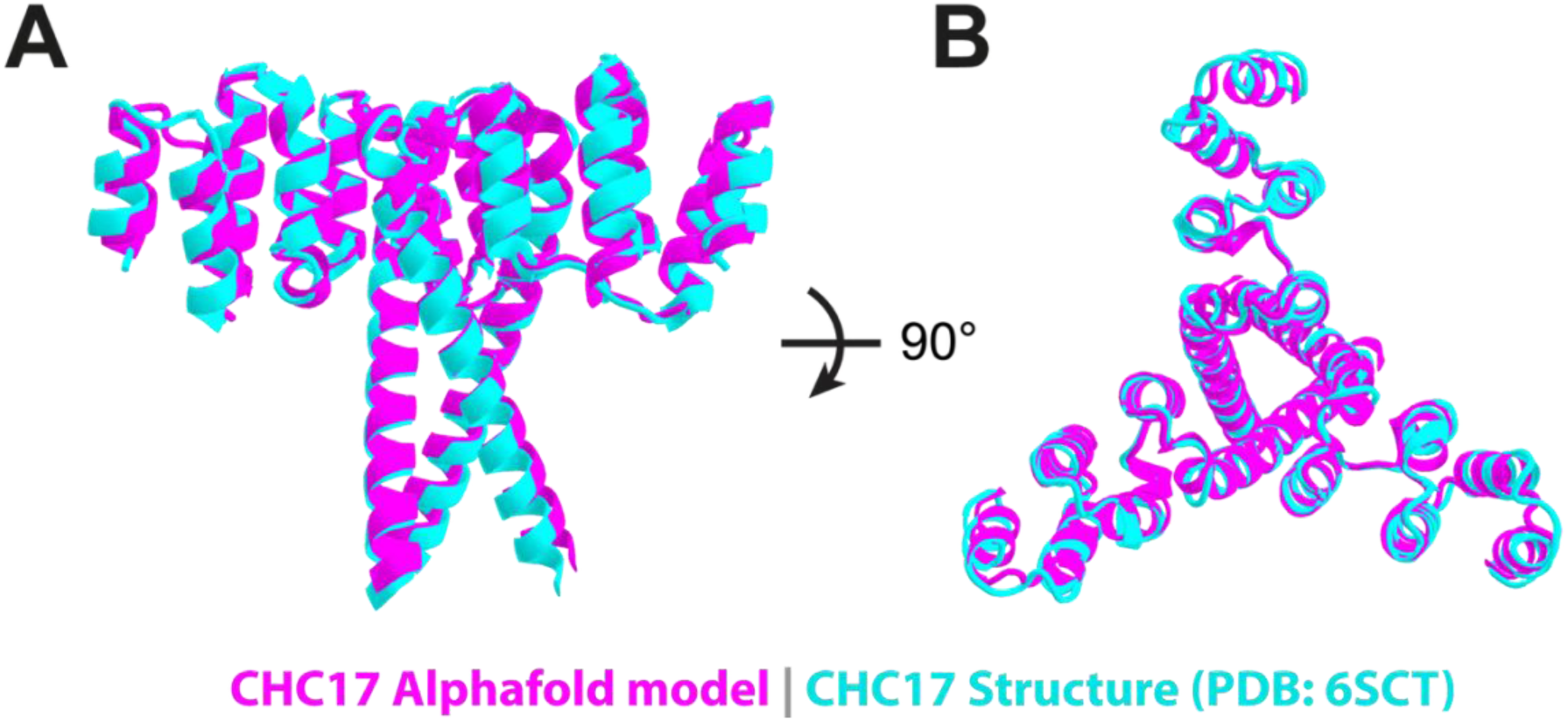
Comparison of CHC17 TxD model and CryoEM structure. Structural alignment of CHC17 TxD (residues 1522 – 1626) generated by AlphaFold-Multimer (magenta) and CryoEM (cyan, PDB: 6SCT) shown from side **(A)** and top views **(B)**.

**Supplementary Figure 2.**
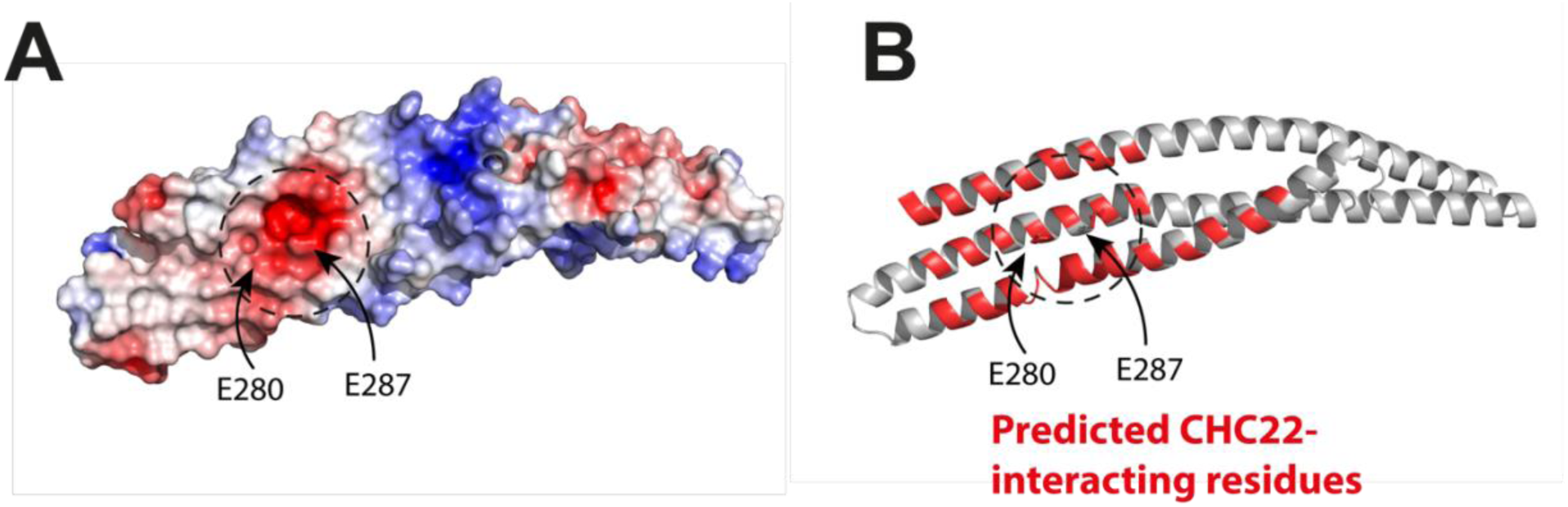
Negative charges of SNX5 may prevent interaction with CHC17 FL. **(A)** The SNX5 BAR domain exhibits a negatively charged hotspot signified by E280 and E287. **(B)** The SNX5 BAR domain is colored based on predicted CHC22 interacting residues (red)^6^. Structure aligned equivalently to **(A)**. The negatively charged hotspot from **(A)** containing E280 and E287 overlaps with the predicted CHC interacting residues.

**Supplementary Table 1 – Clathrin gene locations across 121 mammalian species.**

Table (provided separately as a CSV file) containing species analyzed in Fig. 1, whether CLTCL1 is present and if so the location of the gene.

**Supplementary Table 2.** Buffers used for protein purification, immunoprecipitation and pulldowns.

| Buffer | Composition |
| --- | --- |
| CHC17 IP lysis buffer | 20 mM HEPES pH 7.5, 150 mM NaCl, 1 mM EDTA, 1 mM EGTA, 10% v/v Glycerol, and 0.25% v/v Triton X-100 |
| CHC17Hub binding buffer | 50 mM Tris pH 8, 500 mM NaCl, 25 mM imidazole |
| CHC17Hub elution buffer | 50 mM Tris pH 8, 500 mM NaCl, 500 mM imidazole |
| CHC17Hub SEC buffer | 25 mM Tris pH 8, 150 mM NaCl |
| CLC PD buffer | 50 mM Tris pH 8, 300 mM NaCl, 5% BSA, 0.1% Triton X-100 |
| CLC Wash buffer | 50 mM Tris pH 8, 300 mM NaCl, 0.1% Triton X-100 |
| CHC22TxD binding buffer | 20 mM HEPES pH 7.5, 500 mM NaCl, 20 mM imidazole, 1 mM beta-mercaptoethanol |
| CHC22TxD elution buffer | 20 mM HEPES pH 7.5, 500 mM NaCl, 400 mM imidazole, 1mM beta-mercaptoethanol |
| CHC22 TxD SEC buffer | 20 mM HEPES pH 7.5, 200 mM NaCl, 3 mM beta-mercaptoethanol |
| Native PAGE Cathode Buffer | 50 mM Tris pH 8.9, 70 mM Glycine |
| Native PAGE Anode Buffer | 100 mM Tris pH 7.8 |

**Supplementary Table 3.** CHC22 TxD crystallography data collection and refinement statistics (PDB: 32SP)

|  |  |
| --- | --- |
| **Resolution (Å)** | 46.64 - 2.18 (2.24 - 2.18) |
| **Space group** | I 2 2 2 |
| **Unit cell** | 93.28 112.05 113.88 90 90 90 |
| **Total Number of Unique reflections** | 30947 (2128) |
| **Multiplicity** | 4.2 (3.9) |
| **Completeness (%)** | 93.90 (53.74) |
| **Mean I/sigma(I)** | 10.5 (0.3) |
| **Wilson B-factor** | 31.53 |
| **R-merge (all I+ and I-)** | 0.064 |
| **R-meas (all I+ and I-)** | 0.074 |
| **R-pim (all I+ and I-)** | 0.035 |
| **CC<sub>1/2</sub>** | 0.999 (0.091) |
| **Overall estimate of resolution limit Based on CC<sub>1/2</sub> (Å)** | 2.3 |
| **Reflections used in refinement** | 29403 (1665) |
| **Reflections used for R-free** | 1476 (82) |
| **R-work** | 0.239 (0.815) |
| **R-free** | 0.269 (0.820) |
| **Number of non-hydrogen atoms** | 2877 |
| **macromolecules** | 2780 |
| **ligands** | 74 |
| **solvent** | 19 |
| **Protein residues** | 337 |
| **RMS(bonds)** | 0.008 |
| **RMS(angles)** | 1.54 |
| **Ramachandran favored (%)** | 97.89 |
| **Ramachandran allowed (%)** | 2.11 |
| **Ramachandran outliers (%)** | 0 |
| **Rotamer outliers (%)** | 0.69 |
| **Clashscore** | 7.21 |
| **Average B-factor** | 81.64 |
| **macromolecules** | 82.04 |
| **ligands** | 69.7 |
| **solvent** | 69.95 |
| **Number of TLS groups** | 9 |

*Statistics for the highest-resolution shell are shown in parentheses.*

**Supplementary Table 4 – RNA-seq datasets representing 18 mammalian species.**

Table (provided separately as a CSV file) containing the RNA-seq datasets analyzed in Figure 2, including species, accession codes, tissue and reference genome.

## Author Contribution Statement

G.T.B: conceptualization, investigation, formal analysis, writing - original draft, writing - review and editing. W.P.B: conceptualization, investigation, formal analysis, writing - original draft, writing - review and editing. J.G: conceptualization, investigation, formal analysis, writing - original draft, writing - review and editing. A.M: investigation, formal analysis, writing - review and editing. N.P: investigation, formal analysis, supervision. P.R: investigation. W.S.S: formal analysis, writing - review and editing. K.K: formal analysis, writing - review and editing. R.Z.C: investigation, formal analysis. K.T: supervision, resources, funding acquisition. S.D: formal analysis, writing - review and editing. F.M.B: conceptualization, supervision, resources, funding acquisition, formal analysis, writing - original draft, writing - review and editing.

## Acknowledgements

This work was supported by grants to F.M. Brodsky from the UKRI Medical Research Council (MR/X018377/1), the UKRI Biotechnology and Biological Sciences Research Council (BB/V001221/1) and The Good Food Institute. S. Djordjevic and K. Thalassinos were supported by UKRI Biotechnology and Biological Sciences Research Council grant (BB/V001221/1). Mass spectrometry instruments were funded by Wellcome Trust Multiuser Equipment grant (221521/Z/20/Z) (K. Thalassinos). G.T. Bates was supported by a Wellcome Trust 4-year interdisciplinary PhD studentship (219856/Z/19/Z). We acknowledge the European Synchrotron Radiation Facility (ESRF) for provision of synchrotron radiation facilities.

We thank members of the Brodsky laboratory, past and present, for helpful discussions and support of this project. We additionally acknowledge Letizia Filippi for their assistance with preliminary Native-PAGE experiments.

The cell culture studies reported here use the HeLa cell line. The authors would like to acknowledge Henrietta Lacks, and the HeLa cell line that was established from her tumor cells, which have made significant contributions to scientific progress and advances in human health.

## Data availability statement

The RNA-seq datasets used in this study are available from the Sequence Read Archive (SRA, https://www.ncbi.nlm.nih.gov/sra) and are detailed in Supplementary Table 4. Atomic structure coordinates and crystallographic data are available in the Protein Data Bank (accession code 32SP). The python script for immunofluorescence colocalization analysis in Napari is available on GitHub (https://github.com/kkamuda/napari_2d_colocalisation) and in archived form on Zenodo (https://doi.org/10.5281/zenodo.21775844). All other data is available upon request to the corresponding author.

## References

1. Ferrannini E, Simonson DC, Katz LD, et al. The disposal of an oral glucose load in patients with non-insulin-dependent diabetes. Metabolism. 1988;37(1):79–85.

2. DeFronzo RA, Tripathy D. Skeletal muscle insulin resistance is the primary defect in type 2 diabetes. Diabetes Care. 2009;32 Suppl 2:S157–163.

3. Gould GW, Brodsky FM, Bryant NJ. Building GLUT4 Vesicles: CHC22 Clathrin’s Human Touch. Trends Cell Biol. 2020;30(9):705–719.

4. Camus SM, Camus MD, Figueras-Novoa C, et al. CHC22 clathrin mediates traffic from early secretory compartments for human GLUT4 pathway biogenesis. J Cell Biol. 2020;219(1).

5. Vassilopoulos S, Esk C, Hoshino S, et al. A role for the CHC22 clathrin heavy-chain isoform in human glucose metabolism. Science. 2009;324(5931):1192–1196.

6. Greig J, Bates GT, Yin DI, et al. CHC22 clathrin recruitment to the early secretory pathway requires two-site interaction with SNX5 and p115. EMBO J. 2024;43(19):4298–4323.

7. Esk C, Chen CY, Johannes L, Brodsky FM. The clathrin heavy chain isoform CHC22 functions in a novel endosomal sorting step. J Cell Biol. 2010;188(1):131–144.

8. Dannhauser PN, Camus SM, Sakamoto K, et al. CHC22 and CHC17 clathrins have distinct biochemical properties and display differential regulation and function. J Biol Chem. 2017;292(51):20834–20844.

9. Wakeham DE, Abi-Rached L, Towler MC, Wilbur JD, Parham P, Brodsky FM. Clathrin heavy and light chain isoforms originated by independent mechanisms of gene duplication during chordate evolution. Proc Natl Acad Sci U S A. 2005;102(20):7209–7214.

10. Fumagalli M, Camus SM, Diekmann Y, et al. Genetic diversity of CHC22 clathrin impacts its function in glucose metabolism. Elife. 2019;8.

11. Smith SM, Baker M, Halebian M, Smith CJ. Weak Molecular Interactions in Clathrin-Mediated Endocytosis. Front Mol Biosci. 2017;4:72.

12. Hosaka T, Brooks CC, Presman E, et al. p115 Interacts with the GLUT4 vesicle protein, IRAP, and plays a critical role in insulin-stimulated GLUT4 translocation. Mol Biol Cell. 2005;16(6):2882–2890.

13. Watson RT, Khan AH, Furukawa M, et al. Entry of newly synthesized GLUT4 into the insulin-responsive storage compartment is GGA dependent. EMBO J. 2004;23(10):2059–2070.

14. Lamb CA, McCann RK, Stockli J, James DE, Bryant NJ. Insulin-regulated trafficking of GLUT4 requires ubiquitination. Traffic. 2010;11(11):1445–1454.

15. Blue RE, Curry EG, Engels NM, Lee EY, Giudice J. How alternative splicing affects membrane-trafficking dynamics. J Cell Sci. 2018;131(10).

16. Redlingshöfer L, McLeod F, Chen Y, et al. Clathrin light chain diversity regulates membrane deformation in vitro and synaptic vesicle formation in vivo. Proc Natl Acad Sci U S A. 2020;117(38):23527–23538.

17. Moulay G, Laine J, Lemaitre M, et al. Alternative splicing of clathrin heavy chain contributes to the switch from coated pits to plaques. J Cell Biol. 2020;219(9).

18. Ybe JA, Perez-Miller S, Niu Q, Coates DA, Drazer MW, Clegg ME. Light chain C-terminal region reinforces the stability of clathrin heavy chain trimers. Traffic. 2007;8(8):1101–1110.

19. Ungewickell E. The 70-kd mammalian heat shock proteins are structurally and functionally related to the uncoating protein that releases clathrin triskelia from coated vesicles. EMBO J. 1985;4(13A):3385–3391.

20. Ungewickell E, Ungewickell H, Holstein SE, et al. Role of auxilin in uncoating clathrin-coated vesicles. Nature. 1995;378(6557):632–635.

21. Rapoport I, Boll W, Yu A, Bocking T, Kirchhausen T. A motif in the clathrin heavy chain required for the Hsc70/auxilin uncoating reaction. Mol Biol Cell. 2008;19(1):405–413.

22. Haga Y, Ishii K, Suzuki T. N-glycosylation is critical for the stability and intracellular trafficking of glucose transporter GLUT4. J Biol Chem. 2011;286(36):31320–31327.

23. Evans R, O’Neill M, Pritzel A, et al. Protein complex prediction with AlphaFold-Multimer. bioRxiv. 2022:2021.2010.2004.463034.

24. Jumper J, Evans R, Pritzel A, et al. Highly accurate protein structure prediction with AlphaFold. Nature. 2021;596(7873):583–589.

25. Wilbur JD, Hwang PK, Ybe JA, et al. Conformation switching of clathrin light chain regulates clathrin lattice assembly. Dev Cell. 2010;18(5):841–848.

26. Ybe JA, Fontaine SN, Stone T, Nix J, Lin X, Mishra S. Nuclear localization of clathrin involves a labile helix outside the trimerization domain. FEBS Lett. 2013;587(2):142–149.

27. Morris KL, Jones JR, Halebian M, et al. Cryo-EM of multiple cage architectures reveals a universal mode of clathrin self-assembly. Nat Struct Mol Biol. 2019;26(10):890–898.

28. Towler MC, Gleeson PA, Hoshino S, et al. Clathrin isoform CHC22, a component of neuromuscular and myotendinous junctions, binds sorting nexin 5 and has increased expression during myogenesis and muscle regeneration. Mol Biol Cell. 2004;15(7):3181–3195.

29. Briant K, Redlingshöfer L, Brodsky FM. Clathrin’s life beyond 40: Connecting biochemistry with physiology and disease. Curr Opin Cell Biol. 2020;65:141–149.

30. Zhang Y, Skolnick J. Scoring function for automated assessment of protein structure template quality. Proteins. 2004;57(4):702–710.

31. Trefely S, Khoo PS, Krycer JR, et al. Kinome Screen Identifies PFKFB3 and Glucose Metabolism as Important Regulators of the Insulin/Insulin-like Growth Factor (IGF)-1 Signaling Pathway. J Biol Chem. 2015;290(43):25834–25846.

32. Mantica F, Irimia M. Gene Duplication and Alternative Splicing as Evolutionary Drivers of Proteome Specialization. Bioessays. 2025;47(5):e202400202.

33. Fenton M, Gregory E, Daughdrill G. Protein disorder and autoinhibition: The role of multivalency and effective concentration. Curr Opin Struct Biol. 2023;83:102705.

34. Dandage R, Papkov M, Greco BM, et al. Single-cell imaging of protein dynamics of paralogs reveals sources of gene retention. iScience. 2025;28(7):112771.

35. Nitschke L, Hu RC, Miller AN, Lucas L, Cooper TA. Alternative splicing mediates the compensatory upregulation of MBNL2 upon MBNL1 loss-of-function. Nucleic Acids Res. 2023;51(3):1245–1259.

36. Su Z, Wang J, Yu J, Huang X, Gu X. Evolution of alternative splicing after gene duplication. Genome Res. 2006;16(2):182–189.

37. Teufel AI, Johnson MM, Laurent JM, Kachroo AH, Marcotte EM, Wilke CO. The Many Nuanced Evolutionary Consequences of Duplicated Genes. Mol Biol Evol. 2019;36(2):304–314.

38. Gertz EM, Yu YK, Agarwala R, Schaffer AA, Altschul SF. Composition-based statistics and translated nucleotide searches: improving the TBLASTN module of BLAST. BMC Biol. 2006;4:41.

39. Tamura K, Stecher G, Kumar S. MEGA11: Molecular Evolutionary Genetics Analysis Version 11. Mol Biol Evol. 2021;38(7):3022–3027.

40. Bolger AM, Lohse M, Usadel B. Trimmomatic: a flexible trimmer for Illumina sequence data. Bioinformatics. 2014;30(15):2114–2120.

41. Kim D, Paggi JM, Park C, Bennett C, Salzberg SL. Graph-based genome alignment and genotyping with HISAT2 and HISAT-genotype. Nat Biotechnol. 2019;37(8):907–915.

42. Liao Y, Smyth GK, Shi W. The R package Rsubread is easier, faster, cheaper and better for alignment and quantification of RNA sequencing reads. Nucleic Acids Res. 2019;47(8):e47.

43. Schindelin J, Arganda-Carreras I, Frise E, et al. Fiji: an open-source platform for biological-image analysis. Nat Methods. 2012;9(7):676–682.

44. Chen Y, Briant K, Camus MD, Brodsky FM. Clathrin light chains CLCa and CLCb have non-redundant roles in epithelial lumen formation. Life Sci Alliance. 2024;7(1).

45. Kong AT, Leprevost FV, Avtonomov DM, Mellacheruvu D, Nesvizhskii AI. MSFragger: ultrafast and comprehensive peptide identification in mass spectrometry-based proteomics. Nat Methods. 2017;14(5):513–520.

46. da Veiga Leprevost F, Haynes SE, Avtonomov DM, et al. Philosopher: a versatile toolkit for shotgun proteomics data analysis. Nat Methods. 2020;17(9):869–870.

47. Yu F, Haynes SE, Nesvizhskii AI. IonQuant Enables Accurate and Sensitive Label-Free Quantification With FDR-Controlled Match-Between-Runs. Mol Cell Proteomics. 2021;20:100077.

48. Keegan RM, Winn MD. MrBUMP: an automated pipeline for molecular replacement. Acta Crystallogr D Biol Crystallogr. 2008;64(Pt 1):119–124.

49. Emsley P, Cowtan K. Coot: model-building tools for molecular graphics. Acta Crystallogr D Biol Crystallogr. 2004;60(Pt 12 Pt 1):2126–2132.

50. Murshudov GN, Skubak P, Lebedev AA, et al. REFMAC5 for the refinement of macromolecular crystal structures. Acta Crystallogr D Biol Crystallogr. 2011;67(Pt 4):355–367.

51. Afonine PV, Grosse-Kunstleve RW, Echols N, et al. Towards automated crystallographic structure refinement with phenix.refine. Acta Crystallogr D Biol Crystallogr. 2012;68(Pt 4):352–367.

52. Leaver-Fay A, Tyka M, Lewis SM, et al. ROSETTA3: an object-oriented software suite for the simulation and design of macromolecules. Methods Enzymol. 2011;487:545–574.

53. Stranges PB, Kuhlman B. A comparison of successful and failed protein interface designs highlights the challenges of designing buried hydrogen bonds. Protein Sci. 2013;22(1):74–82.

54. Baker NA, Sept D, Joseph S, Holst MJ, McCammon JA. Electrostatics of nanosystems: application to microtubules and the ribosome. Proc Natl Acad Sci U S A. 2001;98(18):10037–10041.

55. Dolinsky TJ, Nielsen JE, McCammon JA, Baker NA. PDB2PQR: an automated pipeline for the setup of Poisson-Boltzmann electrostatics calculations. Nucleic Acids Res. 2004;32(Web Server issue):W665–667.

56. Mukherjee S, Zhang Y. MM-align: a quick algorithm for aligning multiple-chain protein complex structures using iterative dynamic programming. Nucleic Acids Res. 2009;37(11):e83.

